# RoCK and ROI: Single-cell transcriptomics with multiplexed enrichment of selected transcripts and region-specific sequencing

**DOI:** 10.1101/2024.05.18.594120

**Authors:** Giulia Moro, Izaskun Mallona, Joël Maillard, Michael David Brügger, Hassan Fazilaty, Quentin Szabo, Tomas Valenta, Kristina Handler, Fiona Kerlin, Andreas E. Moor, Robert Zinzen, Mark D. Robinson, Erich Brunner, Konrad Basler

## Abstract

Various tools have been developed to reliably identify, trace and analyze single cells in complex tissues. In recent years, these technologies have been combined with transcriptomic profiling approaches to explore molecular mechanisms that drive development, health, and disease. However, current methods still fall short of profiling single cell transcriptomes comprehensively, with one major challenge being high non-detection rates of specific transcripts and transcript regions. Such information is often crucial to understanding the biology of cells or tissues and includes lowly expressed transcripts, sequence variations and exon junctions. Here, we developed a scRNAseq workflow, RoCK and ROI (Robust Capture of Key transcripts and Regions Of Interest), that tackles these limitations. RoCKseq uses targeted capture to enrich for key transcripts, thereby supporting the detection and identification of cell types and complex phenotypes in scRNAseq experiments. ROIseq directs a subset of reads to a specific region of interest via selective priming to ensure detection. Importantly, RoCK and ROI guarantees efficient retrieval of specific sequence information without compromising overall single cell transcriptome information and our workflow is supported by a novel bioinformatics pipeline to analyze the multimodal information. RoCK and ROI represents a significant enhancement over non-targeted single cell sequencing, particularly when cell categorization depends on transcripts that are missed in standard scRNAseq experiments. In addition, it also allows exploration of biological questions that require assessment of specific sequence elements along the targets to be addressed.

## Main

Single cell RNA sequencing (scRNAseq) is a valuable tool to study gene expression in complex and heterogeneous tissues. Main advances that followed the advent of scRNAseq^1^ are the ability to barcode RNA from individual cells^2^ and the use of barcoded beads to simultaneously analyze many cells^3^. The beads harbor oligonucleotides (oligos) that are covalently attached to and unique to each bead. Through the combination of a single cell with a uniquely barcoded bead in small reaction chambers, the transcripts of a cell can be captured, discernibly barcoded, individually marked (with unique molecular identifiers, UMIs) and processed into cDNA libraries that are suitable for high throughput sequencing (HTS)^3,4^. These advances have fueled the development of various scRNAseq technologies that allow in-depth transcriptional profiling of selected cell populations and acquisition of multimodal datasets for tissue-derived cell mixtures across health and disease^5^.

However, many bead-based high-throughput technologies still suffer from severe limitations. First, the data acquired in such experiments tends to cover a small fraction of each cell’s transcriptome^6–8^. The loss of information may occur at various levels, including mRNA capture on barcoded beads, reverse transcription, preferential PCR amplification during library generation or sequencing bias^9–13^. As a result, a high proportion of expressed genes remain undetected (*i.e.*, a "zero measurement")^8,14,15^. In particular, the detection sensitivity for lowly expressed transcripts remains challenging^8,14^.

Previous methods aiming to mitigate the detection limits of some transcripts can be subdivided into bioinformatic strategies, including handling data as pseudobulks^16,17^, or meta-cells^18^; and wet-lab methods^19–26^. Additional methods, such as targeted amplification^27,28^ require that the transcripts of interest have previously been captured on the beads and reverse transcribed, and therefore do not address the *a priori* problem of capturing rare transcripts in the first place. Similar to targeted amplification, other protocols offer enrichment of transcripts of interest via specific probes at the level of library generation^29,30^ or aim to remove non-informative highly expressed transcripts^31,32^. These methods improve detection of transcripts of interest at the level of the library generation and sequencing, but none addresses the fact that loss of information may already occur during mRNA capture; only targeted capture of transcripts of interest would solve this issue. One solution, DARTseq^33^, used a subset of DNA oligos on barcoded beads that were equipped with nucleotide sequences allowing targeted capture of transcripts of interest. A variable bead modification rate between 25 and 40% was reached, which is reflected by a similar variation of information in the transcriptome profile. Importantly, DARTseq allows both the recovery of transcripts of interest as well as profiles for the transcriptome of cells.

An additional limitation of many scRNAseq methods is the bias toward 3′ or 5′ readouts of dT-captured transcripts^3,34^ resulting in a lower coverage of other regions within the coding sequence (CDS). However, there is often important information in the CDS of a transcript that, if read out, could enhance the value of scRNAseq experiments. This information may be restricted to short regions of interest (ROIs) in a transcript as is the case for splice junctions or single nucleotide variants. In both cases, reads would need to encompass small but specific regions that are unlikely to be efficiently captured by end-directed sequencing. One way to obtain the sequence information of a short ROI is to use full-length sequencing methods^19,22,35–37^ or VASAseq^31^, which is based on fragmentation of all RNA molecules in a cell followed by polyadenylation, providing information across the full transcript by other means. Other solutions aiming to detect ROIs in transcripts rely on specific primers or additional amplification steps^20,26,38–41^, all of which significantly increase complexity of library generation protocols. Furthermore, these approaches have in common that they use pre-amplified cDNA or synthesized first strands of dT-captured transcripts to amplify the ROI.

Although the described methods increase the detection of transcripts and tackle the technical challenges of sequencing through a ROI, they also come with limitations such as lengthy library generation protocols, increased sequencing cost or the inability to target multiple regions of interest. To target regions and transcripts of interest as well as profiling the full transcriptome of cells, we developed RoCK and ROI, a novel and simple scRNAseq workflow. The method combines targeted capture, termed <u>RoCK</u>seq (Robust Capture of Key transcripts), with <u>ROI</u>seq (Region Of Interest), for specific detection of features of interest by HTS. Importantly, a standard whole transcriptome analysis (WTA) library is generated for the same cells without sacrificing detection depth and in a manner that allows cell-by-cell pairing of WTA, RoCKseq and ROIseq information. RoCK and ROI can achieve robust target detection in up to 98% of cells. By validating RoCK and ROI in multiple biological samples, we show that we can complement the transcriptome of cells with the sequence information of specifically captured transcripts and reads directed to regions of interest in the same workflow. We anticipate that RoCK and ROI will be widely applicable across biological samples, significantly improve detection of crucial features and allow new insights into biological mechanisms at the single cell level.

## Results

### RoCKseq bead modification is reproducible, long-lasting and titratable

In order to detect specific mRNAs in single cells, we established a method that captures mRNAs not via the polyA tail but rather through hybridization to an upstream target site such as within the coding sequence (CDS; see Figure 1a, Supp Figure 1a). The method uses barcoded beads commercially available for the BD Rhapsody scRNAseq platform, for which the sequence information of the bead-attached oligos is publically accessible (Supp Figure 1b). The beads carry two types of oligos: i) dT oligos, which are needed to capture polyadenylated mRNAs to obtain WTA information; and ii) template switching oligos (TSO), standardly used for the VDJ full length (TCR/BCR) assay. To specifically capture mRNAs of interest, we reasoned that it should be possible to append the TSO oligos with a capture sequence complementary to the target(s) of interest (referred to as RoCKseq beads). Additionally, by only modifying the TSO-portion we would not compromise the beads’ ability to generate WTA libraries. To establish the method, we first focused on appending a single capture sequence complementary to the *eGFP* CDS to the TSOs (Supp Figure 2a-b). The addition of the capture sequence is mediated by a DNA polymerase-based enzymatic reaction using a single stranded DNA oligo (splint) for modification (Figure 1b, Supp Figure 2a-b). After annealing of the splint to the TSOs, the T4 DNA polymerase elongates the recessed ends, generating double stranded DNA molecules. Since the T4 DNA polymerase has an intrinsic 3’-5’ exonuclease activity that targets single-stranded DNA (ssDNA)^42,43^, a phosphorylated polyA oligo is added to protect the dT oligos at this step. Importantly, to restore the modified TSO and dT oligos to the single-strandedness needed for mRNA capture, we use a lambda exonuclease to remove the complementary strand. This enzyme strongly prefers phosphorylated 5’-ends compared to unphosphorylated DNA^44,45^, hence the addition of the 5’ phosphate groups to the splint and protective oligos.

**Figure 1:**
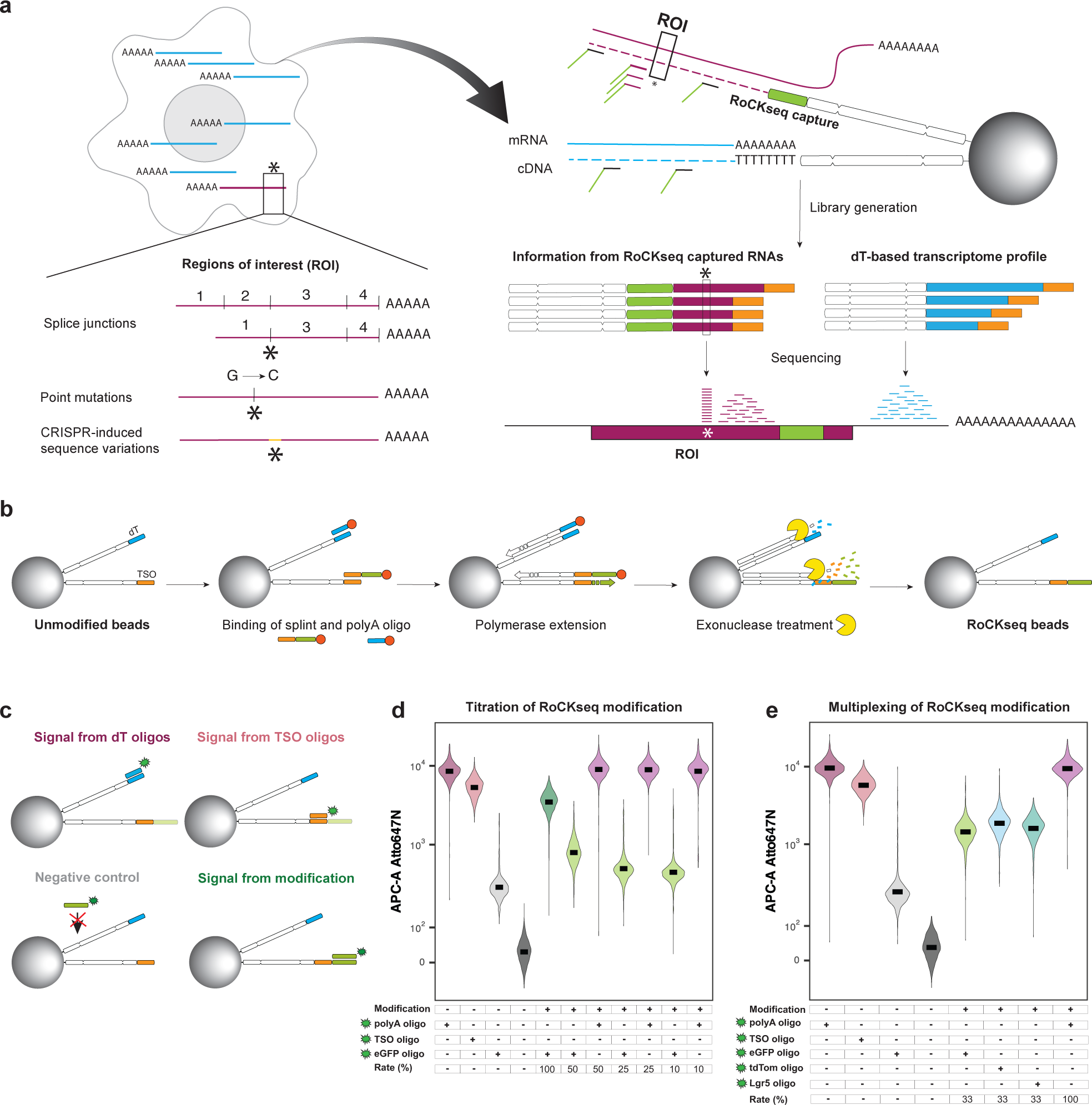
RoCK and ROI concept and examples of RoCKseq bead modification. **a,** Technique overview including BD Rhapsody beads modification (RoCKseq) and regions of interest enrichment (ROI) via primer addition. **b**, RoCKseq bead modification. Red circles: 5’ phosphate groups on oligonucleotides. **c,** Fluorescent assay to assess bead modification and quality of oligos on the beads. Signal from dT oligos: polyA probe binding to the dT stretch on the beads; Signal from TSO oligos: probe complementary to the TSO; Negative control: probe complementary to the capture on unmodified beads; Signal from modification: probe complementary to the capture on the modified beads. **d-e,** FACS quantification of RoCKseq bead modification. Titration of modification on RoCKseq beads ranging from 100% to 10% **(d)**. Target: *eGFP* CDS. Modification of RoCKseq beads with multiple capture sequences in the same ratio (33% each) **(e)**. Targets: *eGFP* CDS, *tdTomato* CDS, *Lgr5* CDS. To assess integrity of dT oligos on modified beads and to determine splint removal by lambda exonuclease, beads were tested using a polyA fluorescent oligo. For panels **d-e**, Y-axis: Atto647N fluorescent signal. The Y-axis has a biexponential transformation.

**Figure 2:**
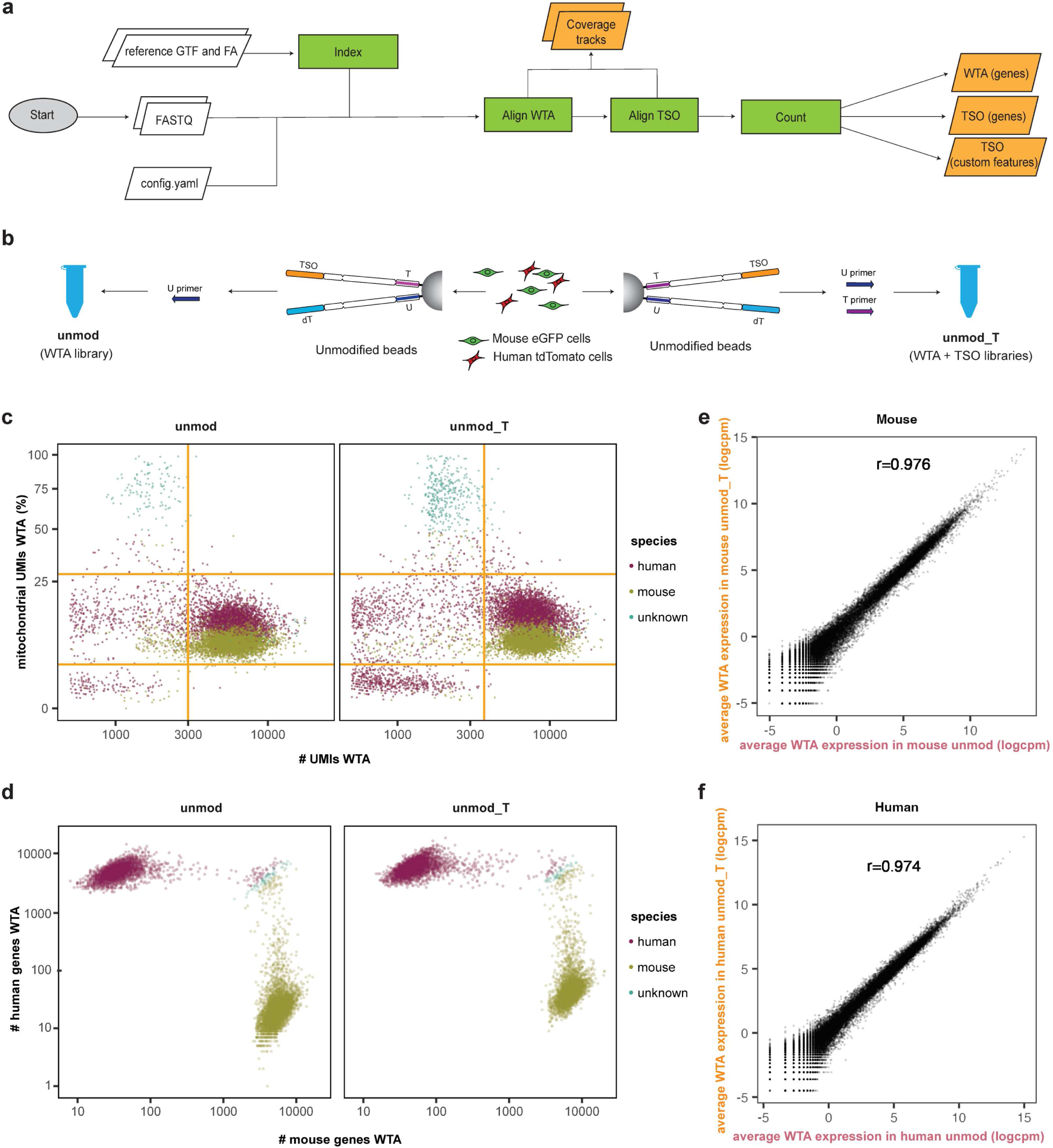
Analysis workflow and testing of the effect of addition of T primer on the WTA of unmodified beads. **a,** RoCK and ROI data analysis pipeline (see Methods). **b,** Experimental setup (mixing scRNAseq experiment) including unmod (U primer) and unmod_T (U and T primers) conditions. **c,** QC of WTA data depicting filtering thresholds (orange). **d,** Barnyard plot depicting cell species assignment using WTA data. **e,** Correlation of WTA readouts in unmod_T versus unmod conditions, mouse cells only. **f,** Same as **(e)** for human cells.

To assess the extent of RoCKseq modification, a fluorescent assay that tests the binding capacity of distinct fluorescent probes was implemented (Figure 1c; see also Saikia et al, 2019^33^). Using this assay, we verified that various modification rates can be easily obtained using the RoCKseq bead modification protocol (Figure 1d). This is achieved using a mixture of the splint and an oligo that is complementary to the TSO sequence but lacks the capture sequence (TSO titration oligo, Supp Figure 1c). In addition, this experiment shows that the bead modification does not alter bead integrity (including dT oligos; Figure 1d) or size (Supp Figure 1d-f). Furthermore, we successfully validated multiplexed modification with three splints (Figure 1e). The importance and efficiency of the lambda exonuclease step was tested by comparing the standard treatment with a sample where either the entire step or the addition of the enzyme was omitted (Supp Figure 3a). We observed that incubation with lambda exonuclease was necessary to fully restore the single strands on the beads (lower fluorescent signal for the other two conditions). In a next step, the effect of the protective polyA oligos used to prevent degradation of dT oligos by the T4 polymerase on the beads was tested. As shown in Supp Figure 3b, both the dT and TSO oligos were degraded if they remained unprotected. The addition of the protective TSO oligo during bead modification is thus important to keep into consideration when modification rates are lower than 100%.

**Figure 3:**
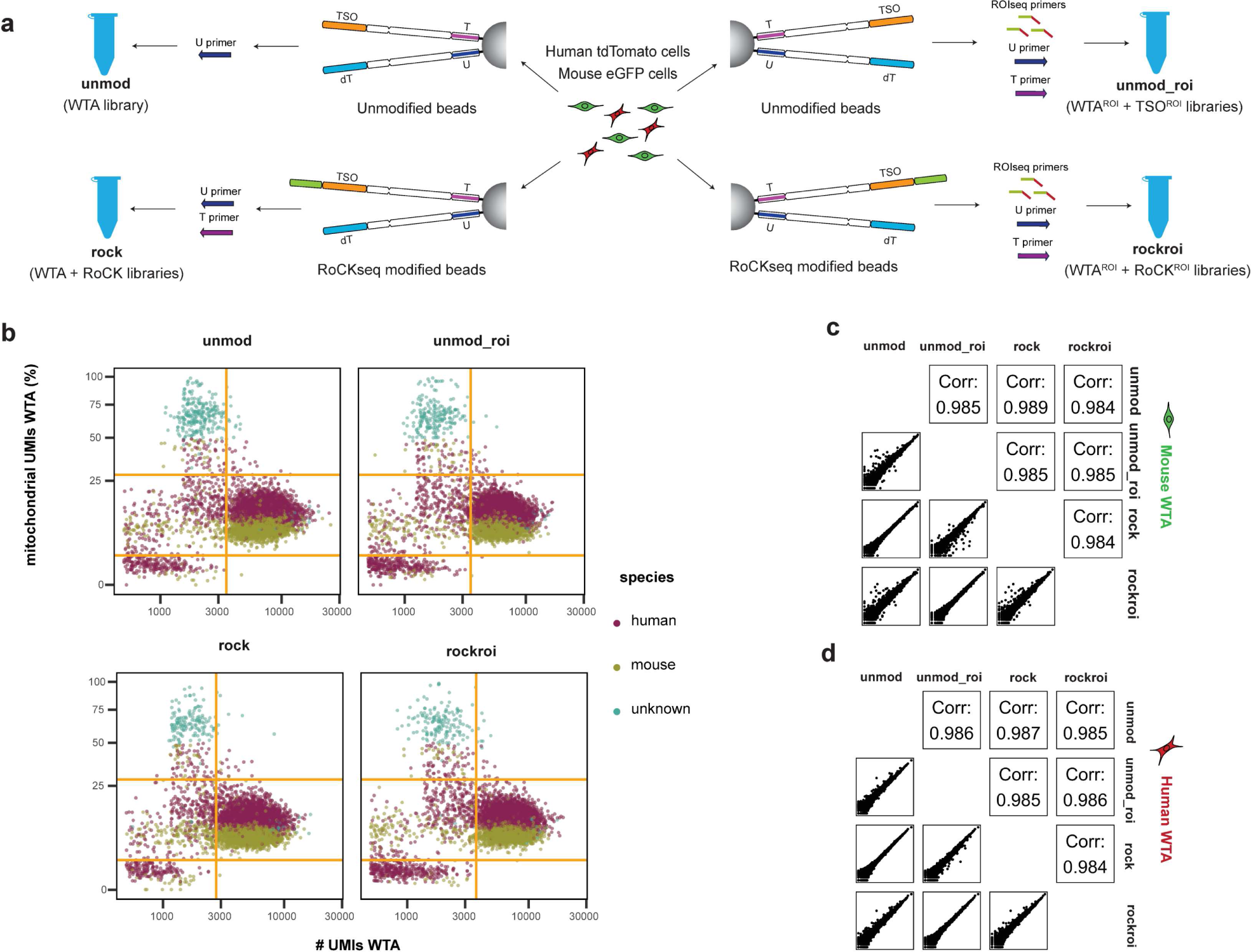
Analysis of RoCK and ROI WTA data from mixing experiment. **a,** Experimental setup of the mixing experiment with extended conditions, including unmodified beads (unmod, unmod_roi) and modified beads (rock, rockroi). Primers: U (all conditions); T: unmod_roi, rock, rockroi; ROIseq: unmod_roi and rockroi conditions. **b,** QC of WTA data depicting filtering thresholds (orange). **c,** Pairwise correlation of WTA data across conditions (mouse cells only)**. d,** same as **(c)** for human cells.

To optimize the modification protocol, various other parameters were tested, including preincubation of the beads with the splint/polyA mix, prewarming of splints (Supp Figure 3c-d) and purification level of oligos (Supp Figure 3e). Importantly, we observe that RoCKseq modification is highly reproducible (Supp Figure 3f) and modified beads remain stable over extended periods of time (at least 19 months; Supp Figure 3g).

Taken together, these results show that standard BD Rhapsody barcoded beads can be reproducibly modified with custom capture sequences while maintaining bead integrity and with low variation among the pool of modified beads. Furthermore, distinct modifications (multiplexed capture sequences) can be easily combined and the rate of modification is scalable, producing custom RoCKseq beads that remain stable for months.

### Reads are directed to regions of interest using ROIseq

To direct reads to regions of interest, we developed ROIseq, in which a specific primer (or multiple primers for multiple ROIs) is (are) spiked into the pool of randomers during library generation (Supp Figure 4a-b). Randomers are random primers of nine nucleotides to which an adapter is attached, and which are used to generate cDNA second strands in the BD Rhapsody platform. Importantly, the addition of randomers leads to the generation of random 5’ ends for the cDNAs (Supp Figure 4a). By specifically designing primers targeting regions of interest (ROIs) on target mRNAs, we can enrich for pre-defined 5’ ends of the corresponding cDNAs and thus specifically guide the reads obtained by HTS-based analysis to the ROIs in the target transcript (Supp Figure 4b). Importantly, the standard randomers used for library generation are also included to profile the cell’s transcriptome. To obtain information on both the transcriptome of a cell and the targeted capture library in the same experiment, a novel library generation protocol was developed (Supp Figure 4b). The new library entails four main changes to the standard BD Rhapsody library generation protocol (Supp Figure 4a). First, a T primer (specific to TSO oligos on the BD Rhapsody beads; see Supp Figure 1a-b) is added during second-strand PCR amplification to retrieve information from the RoCKseq captured transcripts. Second, ROIseq primers are added to the pool of randomers to direct reads to regions of interest. Next, a custom indexing primer is used for the indexing of the RoCKseq capture library. This leads to the generation of two separately indexed libraries, one derived from the dT oligos (WTA library) and the other from TSO oligos (TSO library) that are mixed for HTS sequencing. Finally, a custom primer is added during HTS sequencing to retrieve information from TSO libraries.

**Figure 4:**
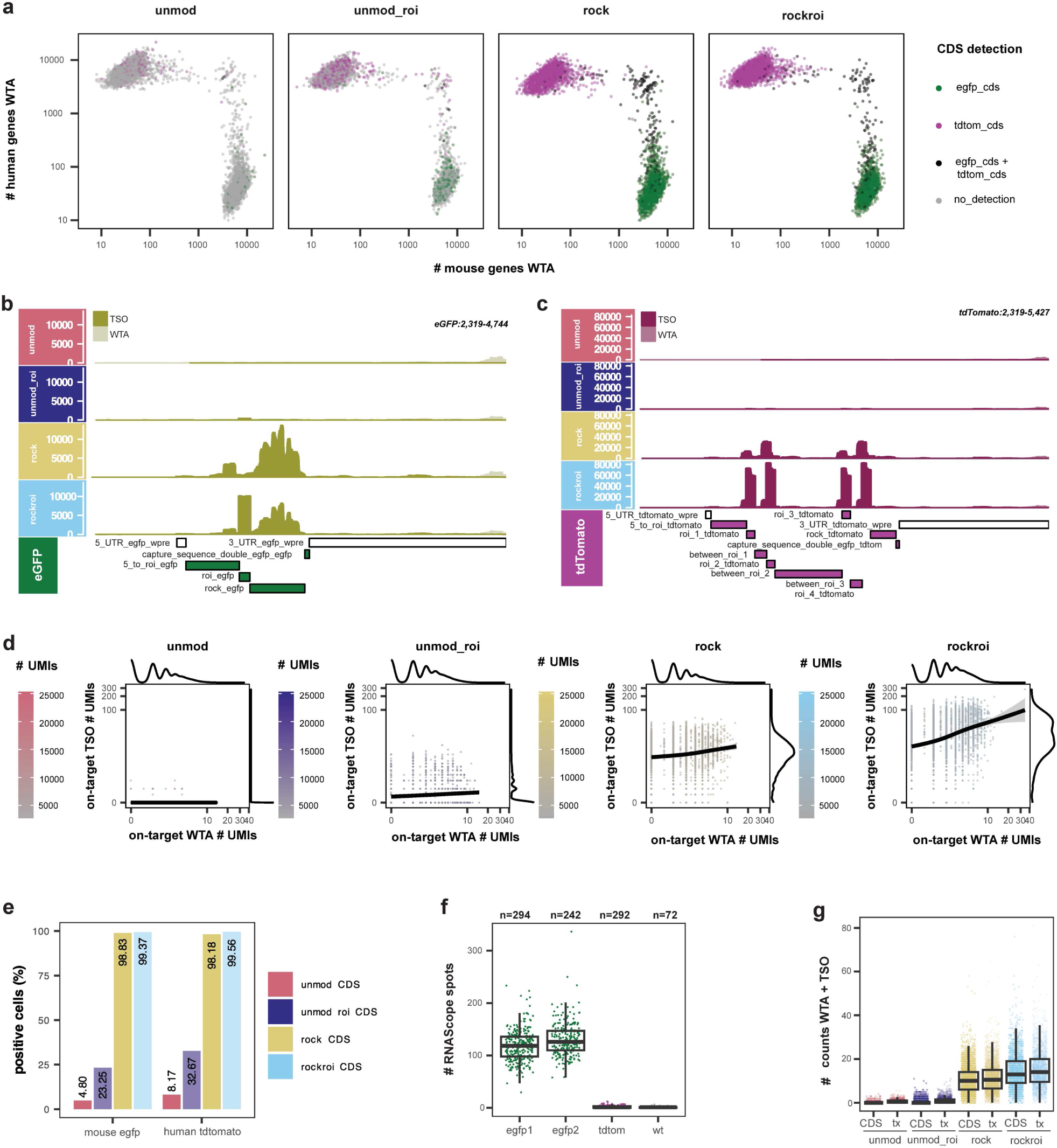
Analysis of RoCK and ROI target enrichment data and quantification of *eGFP* mRNAs. **a,** Barnyard plot colored by detection of *eGFP* and *tdTomato* CDS in WTA and TSO data. **b,** Sequencing coverage and depth along *eGFP* in mouse cells for TSO (olive green) and WTA (off white). **c,** Sequencing coverage and depth along *tdTomato* in human cells for TSO (red purple) and WTA (light mauve). **d**, Detection of *eGFP* and *tdTomato* in TSO versus WTA data, per cell**. e**, Percent of cells with detectable *eGFP* CDS (mouse cells) or *tdTomato* CDS (human cells) in TSO plus WTA data, per condition **f,** Number of *eGFP* mRNAs in mouse cells detected by RNAScope. egfp1 and egfp2: replicates. Negative controls: L-cells expressing tdTomato and wt L-cells (untransduced). **g,** Number of UMIs from combining WTA and TSO data for the *eGFP* CDS and transcript (tx).

### A custom, reproducible and automated workflow to analyze targeted and untargeted data

We have designed an open-source Snakemake^46^ workflow to process data from raw sequencing reads, leveraging the BD Rhapsody dual oligos present on the barcoded beads, with distinctive cell barcode structure differentiating the targeted (TSO) from untargeted (WTA) data (Supp Figure 1b; see Methods). The workflow (Figure 2a) generates a transcriptome index to match the experimental design (*i.e.,* taking into account the cDNA read length). After indexing with STAR^47^, FASTQ files are aligned and counted using STARsolo^48^ while extracting valid cell barcodes and producing count tables for TSO and WTA readouts separately. We provide other running modes to deal with ad-hoc use cases, such as targeting repetitive sequences and hence including multimapping reads. Aside from producing count tables, our workflow generates basic scRNAseq analysis reports, including quality control and cell clustering.

### Addition of T primer during RoCK and ROI library generation does not affect WTA information

To test the RoCK and ROI concept, we wanted to confirm that the addition of the T primer does not affect the WTA readouts. Additionally, since this primer is needed to obtain information from TSO oligos, we aimed to explore the generation of a TSO oligo-based library (TSO library) given by the capture of transcripts on these oligos. For this assessment, we chose two clonal cell lines, each expressing distinct fluorescent proteins. We generated a 1:1 mix of clonal (human) HEK293-T cells expressing tdTomato and clonal (murine) L-cells expressing eGFP, both of which were generated by lentiviral transduction (Supp Figure 5a). We generated libraries using unmodified beads and a standard BD Rhapsody protocol, either with (unmod_T, WTA and TSO libraries) or without (unmod, WTA library) the addition of the T primer (Figure 2b, Supp Table 1). A first evaluation of the libraries before indexing did not reveal any noticeable difference between the two conditions (Supp Figure 5b). We then reasoned that the addition of the T primer would have generated cDNAs from transcripts that have been captured by the TSO sequence (TSO library). We therefore indexed the standard dT libraries (with and without T primer) as well as the TSO library generated from the putative TSO captured mRNAs. As before, the two final dT libraries had very similar characteristics (Supp Figure 5c). As presumed, also a TSO library was generated with a similar trace as the dT libraries. This indicates that the addition of a T primer allows the retrieval of information that derives from transcripts captured via the TSO.

**Figure 5:**
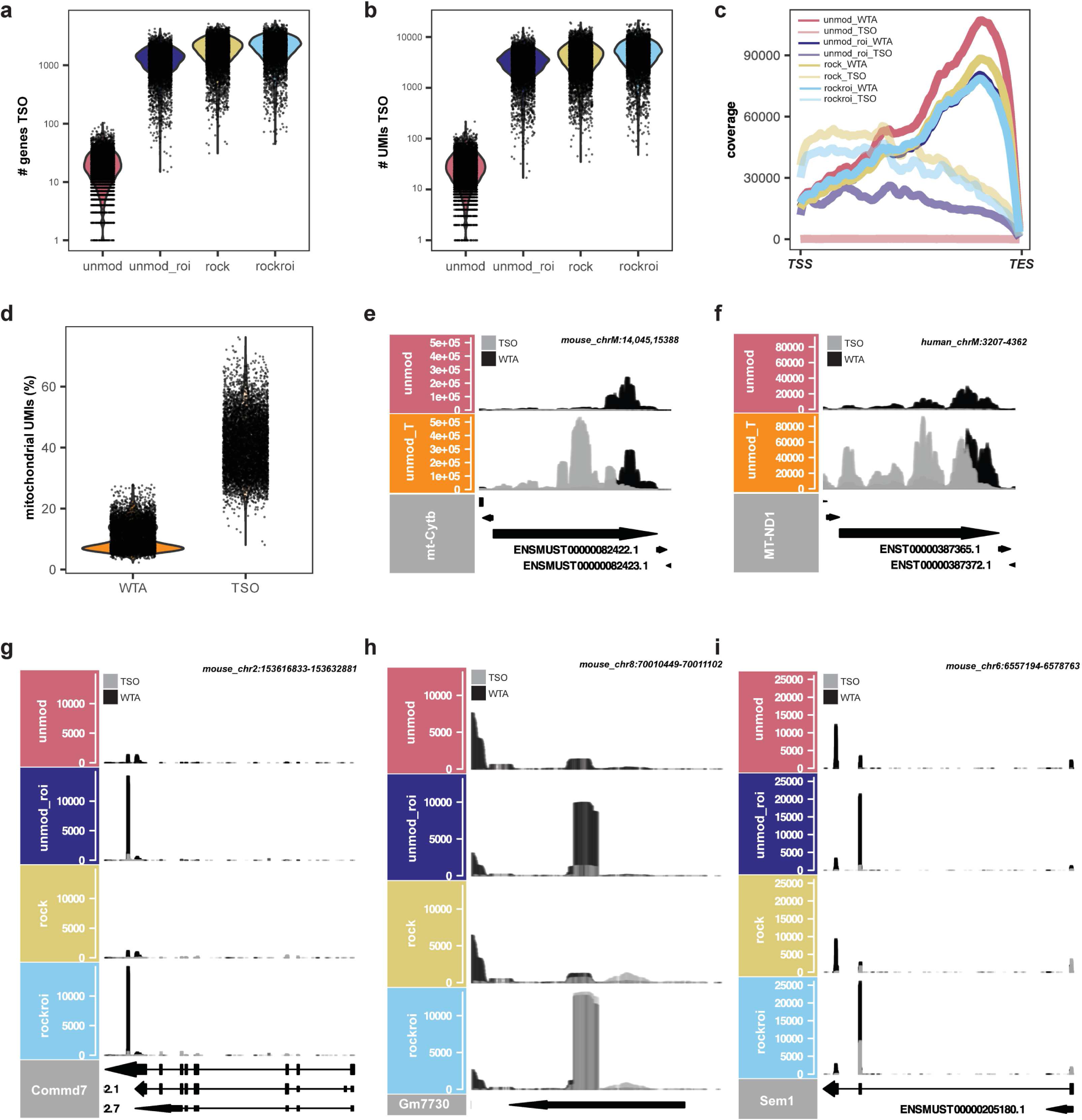
Characterization of RoCK and ROI TSO data and example of ROIseq peaks. **a,** Number of genes detected in the TSO data. **b,** Number of UMIs detected in the TSO data. **c,** Aggregated sequencing coverage along detected transcripts for TSO and WTA data; TSS: transcription start site; TES: transcription end site. **d,** Mitochondrial content in WTA and TSO data. **e-i,** Sequencing coverage for TSO (gray) and WTA (black) along *mt-Cytb* **(e)**, *MT-ND1* **(f)**, *Commd7* **(g)**, *Gm7730* **(h)** and *Sem1* **(i).** Data in panels (**a-c** and **g-i**) refers to experiment described in Figure 3 **(a)**, data in panels **(d-f)** refers to experiment described in Figure 2 **(b)**.

After sequencing and single-cell read mapping and counting, we first checked whether the information from the two WTA libraries was similar in terms of number of genes (Supp Figure 5d), number of UMIs (Figure 2c, Supp Figure 5e) and percent of mitochondrial content (Figure 2c, Supp Figure 5f) detected per cell. This was the case for both human and mouse cells. The two cell types could be clearly distinguished based on the WTA libraries (Figure 2d). To determine if the T primer addition affects the WTA readout, we pairwise compared the per-gene counts obtained with and without addition of the T primer. The transcriptomes of the unmod and unmod_T conditions were similar (Figure 2e-f, Pearson correlation 0.976 for mouse, 0.974 for human), indicating that the T primer does not hamper library generation nor significantly alter the WTA signal derived from dT oligos.

### RoCKseq and RoCK and ROI target transcripts in a sensitive and specific manner and do not bias the WTA information

We next performed a RoCK and ROI experiment using the same cell lines (1:1 mix of eGFP or tdTomato expressing cells). The aim of the experiment was to compare the *eGFP* and *tdTomato* detection sensitivity with and without targeted capture (RoCKseq) and ROIseq-based priming. To capture both transcripts, we selected a 25 bp stretch at the 3’ end of the CDSs that is shared between *eGFP* and *tdTomato* (Supp Figure 5g), allowing a single configuration of RoCKseq beads (Supp Figure 6a). A single ROIseq primer for *eGFP* and two ROIseq primers for *tdTomato* were used. Additionally, both transcripts share the 5’ and 3’ UTR sequences, and hence can only be distinguished by reads from their respective CDSs (Supp Figure 5a).

**Figure 6:**
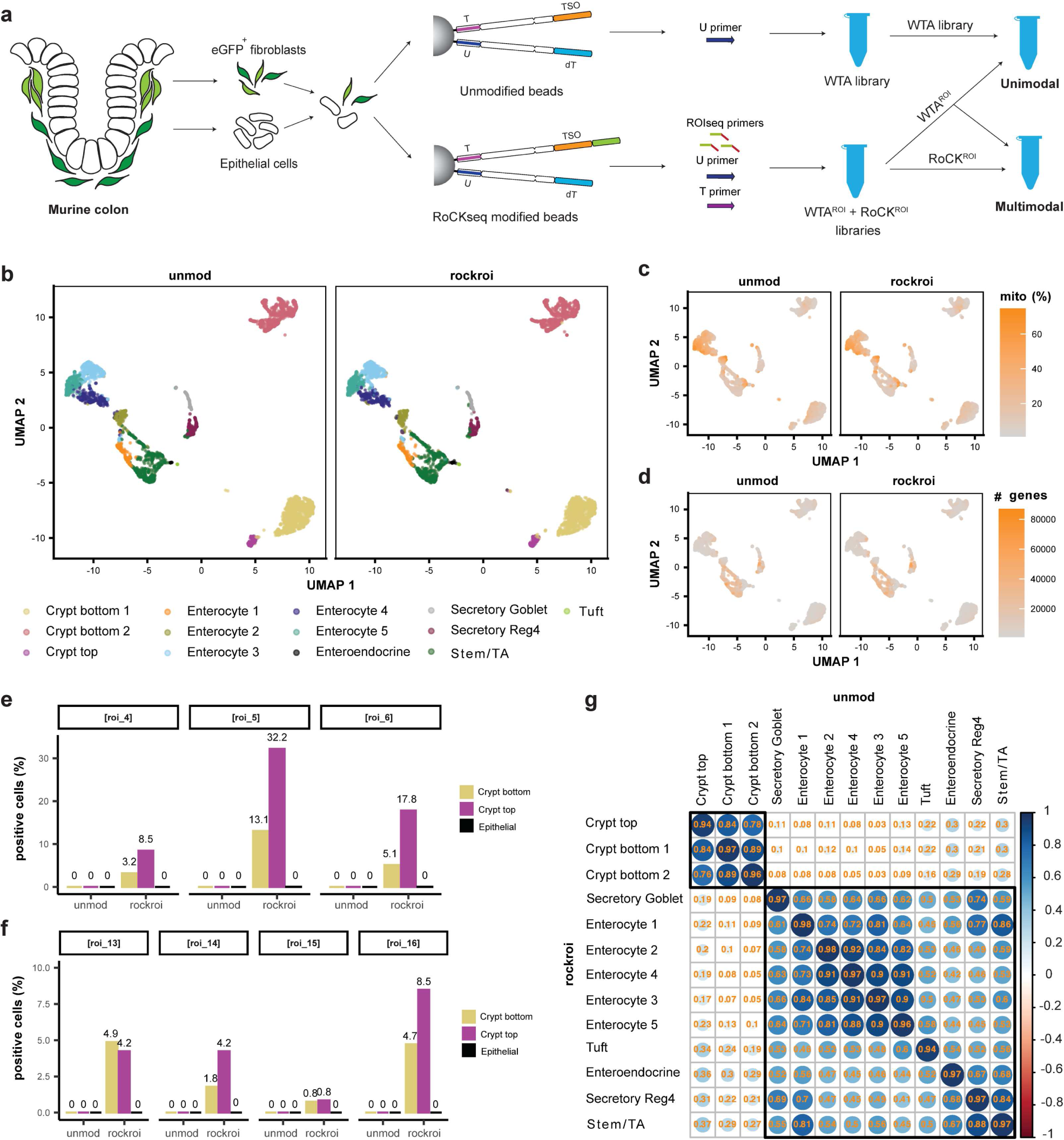
Detection of *Pdgfrα* splice junctions in murine colon cells. **a,** Experimental set up of the *Pdgfrα* experiment including eGFP+ fibroblasts and EpCAM+ epithelial cells from murine colon. Conditions: unmod and rockroi (unimodal and multimodal); Primers: U (all conditions); T and ROIseq (rockroi). **b,** UMAP embedding on unimodal WTA data split by bead modification and colored by cell type (unsupervised clustering). **c-d,** Same UMAP colored by mitochondrial content **(c)** and by number of detected genes in WTA **(d). e-f,** TSO detection rate of ROIseq-targeted splice junctions. **g**, Pairwise Pearson correlation of gene (WTA) expression readouts across cell types in unmod (horizontal) versus rockroi conditions (vertical).

To assess the individual effects of RoCKseq capture and ROIseq primers, we tested four experimental conditions (Figure 3a, Supp Table 1): the standard BD Rhapsody protocol using i) unmodified beads (unmod) or ii) the unmodified beads with ROIseq and T primers (unmod_roi); and RoCKseq beads iii) without (rock) and iv) with the addition of ROIseq primers (rockroi). Initial quality control on libraries before and after indexing (Supp Figure 6b and 6c, respectively) showed that global properties of the WTA and TSO libraries were similar for all conditions.

The four samples had similar WTA transcriptomes in terms of number of genes, number of transcripts and percent mitochondrial content (Figure 3b, Supp Figure 6d-f). In addition, the information in the WTA libraries was sufficient to distinguish between mouse and human cells in all conditions (Supp Figure 6g). The similarity of the average transcriptional profiles across conditions was apparent when comparing the information obtained from the WTAs for mouse (Figure 3c, Pearson correlations between 0.984 and 0.989) and human (Figure 3d, Pearson correlations between 0.984 and 0.987) samples.

We next focused on the CDS detection for the *eGFP* and *tdTomato* transcripts. Compared to the unmod condition, unmod_roi and particularly rock and rockroi showed an increase in the number of cells with at least one detected UMI in the respective *eGFP* and *tdTomato* CDS (Figure 4a). This was particularly apparent in the rock and rockroi conditions, indicating that RoCKseq capture strongly aids with the detection of the CDS. This is also seen when looking at the coverage along the *eGFP* and *tdTomato* transcripts in mouse and human cells, respectively (Figure 4b-c). Compared to rock, rockroi highlights a single prominent peak of reads in the *eGFP* transcript precisely where the ROIseq primer had been positioned. Similarly, two distinctive coverage signal peaks can be seen in the rockroi condition for the *tdTomato* transcript. Since *tdTomato* was generated by fusing two copies of the dTomato gene to create a tandem dimer^49^, we retained multimapping alignments, hence reporting alignments twice. Of note, when comparing the unmod_roi with the rockroi condition, the need for RoCKseq capture when targeting sequences of interest becomes apparent, as only a small peak of reads is visible in the unmod_roi condition. This can be explained by the distance of the CDS to the polyA tail being >1.5 kb in all cases. The WTA coverage remained very similar across conditions (Supp Figure 7a-b). The increase in sequencing coverage of the *eGFP* and *tdTomato* CDSs is largely driven by TSO reads (Figure 4b-c). This is also apparent when comparing the numbers of UMIs per cell derived from WTA versus TSO (Figure 4d).

We next looked at the percent of cells with detectable *eGFP* and *tdTomato*. Due to the *eGFP* and *tdTomato* sequence similarity we consider as positive cells those with at least one alignment to the targeted CDSs, unique or not. Compared to the unmod condition, where *eGFP* CDS and *tdTomato* CDS was detected in 4.80% and 8.17% of cells, respectively, RoCKseq capture increased the detection to 98.83% *eGFP-*positive L-cells and 98.18% *tdTomato*-positive HEK293-T cells (Figure 4e). The addition of the ROIseq primer (unmod_roi) increased the detection of the *eGFP* CDS and *tdTomato* CDS to 23.25% and to 32.67%, respectively, while in rockroi an even higher proportion of positive cells was detected (*eGFP* CDS: 99.37%; *tdTomato* CDS: 99.56%). The detection of *eGFP* and *tdTomato* was highly specific, with very low false positives for the RoCKseq and ROIseq regions (for mouse cells: Supp Figure 7c, for human cells: Supp Figure 7d).

To evaluate the sensitivity of RoCK and ROI, we wanted to understand how the number of UMIs relates to the number of targeted mRNAs present in a cell. Previous reports have shown that only 5-20% of the transcriptome of a cell is recovered in scRNAseq experiments^6–8,50,51^. To determine the number of *eGFP* transcripts expressed in a cell, we visually detected single transcripts by RNAscope on the same clonal L-cell line (Supp Figure 7e-g). As a negative control, we used untransfected L-cells (wt) or a clonal L-cell line expressing tdTomato. For *eGFP* transcripts, we quantified 30-233 spots (average 118, median 118.5) for the first replicate and 58-336 spots (average 131, median 126) for the second replicate, both varying according to cell size (Supp Figure 7h; Supp Figure 7i RNAScope spots normalized by area). On the other hand, an average of 0.52 counts per cell were measured in the scRNAseq experiment for the full *eGFP* transcript (CDS plus UTRs) in the unmod condition, 11.28 counts for the rock condition and 15.29 counts per cell for the rockroi condition (Figure 4f-g). This indicates that we detect 0.42% of *eGFP* transcripts per cell for the unmod condition and 12.30% for the rockroi condition, thus reaching the transcript detection limit indicated in previous reports^6–8,50,51^ with our method.

Altogether, RoCKseq beads lead to a drastic increase in the detection of transcripts of interest and in combination with ROIseq primers, RoCK and ROI enriches reads in regions of interest. This targeted information is recorded together with the WTA of cells.

### Characterization of RoCKseq capture and ROIseq targets

We next looked specifically into the TSO modality for reads that did not map to our targeted regions. For the scRNAseq experiment described in Figure 3a, the percentage of (on-target) *eGFP-* or *tdTomato*-specific TSO alignments was 0.5%, 0.22% and 0.01% for the rockroi, rock and unmod_roi conditions, respectively (Supp Figure 8a), indicating low specificity. This was also apparent when looking at the number of genes and UMIs detected in the TSO data in all samples in which the T primer was added, independent of bead modification (Figure 5a-b) and was also true for the scRNAseq experiment described in Figure 2b (Supp Figure 8b-c). The percentage of intergenic information was slightly higher in WTA compared to TSO libraries (Supp Figure 8d). Additionally, the TSO information in genes showed a higher percentage of non-protein coding genes compared to the WTA libraries (Supp Figure 8e), including also non-polyadenylated types, which may be explained by internal capture of transcripts. In fact, when looking at the TSO coverage across gene bodies, it was not biased towards the transcript 3’ end (as is the WTA readout; Figure 5c). This was true for both scRNAseq experiments and thus independent of the bead modification (Supp Figure 8f). Compared to the WTA readouts, the unmod_T TSO modality showed a higher percentage of mitochondrial transcripts per cell (Figure 5d). This was also apparent when looking at the coverage across the detected mitochondrial transcripts (Figure 5e-f).

Compared to the WTA modality, TSO libraries had a lower number of genes (Supp Figure 6d versus Figure 5a), UMIs (Supp Figure 6e versus Figure 5b) and reads with canonical cell barcodes (Supp Figure 8g), although the libraries were mixed at a 1:1 concentration. The TSO libraries also had a lower number of alignments compared to the WTA libraries (Supp Figure 8h). To look into this, we tracked the reads and alignments across the conditions of the two mixing experiments at different relevant steps for the data analysis (Supp Figure 9a-l). We noticed most WTA aligned reads belonged to high-quality (retained) cell barcodes regardless of the bead modification, whereas most TSO reads did not. This difference is also reflected when looking at the total number of UMIs in the TSO and WTA count tables. The discrepancy in the amount of information deriving from WTA and TSO modalities is thus occurring already at the sequencing step and is further affected by downstream processing steps.

Although we observed this difference in the two data modalities and the percentage of *eGFP-* or *tdTomato*-specific TSO alignments is low, the number of on-target UMIs was higher in TSO versus WTA data in all conditions in which the T primer was added and especially for rock and rockroi; 80-fold and 94-fold for on-target (unique or not) alignments in rock and rockroi respectively (Supp Figure 9j and Supp Figure 9l).

Similar to the RoCKseq capture, we asked if the ROIseq primers bind in transcripts other than the targeted *eGFP* and *tdTomato*. We observed that the ROIseq primers were binding to off-target mRNAs, leading to ROIseq-specific peaks on both WTA and TSO modalities (Figure 5g-i). On the other hand, we found that the WTA in modified beads is very similar to that of unmodified beads (Figure 3c-d), indicating that neither RoCKseq nor ROIseq had a major impact on the overall untargeted transcriptome.

### RoCK and ROI enables the detection of *Pdgfrα* splice junctions in murine colon cells

After validation of RoCK and ROI in cell lines, we chose the murine colon as a complex biological system with multiple transcriptionally-distinct cell types (Figure 6a, Supp Table 1). First, we wanted to test if the WTA modality from a RoCK and ROI experiment can identify and annotate the same cell types as that from an unmodified bead experiment. Second, we wanted to quantify splice junctions of a targeted transcript. We chose a mouse strain where the H2B-eGFP fusion protein reporter construct was knocked into one of the *Pdgfrα* alleles^52^ (Supp Figure 10a-b), where *Pdgfrα* is a marker for mesenchymal cells. Of note, the *Pdgfrα* gene (and the *eGFP* reporter) is expressed at different levels in crypt top and crypt bottom fibroblasts^53^. Several protein-coding transcripts are encoded by the wildtype *Pdgfrα*, including short transcripts with 16 exons and long transcripts with seven additional exons. For RoCK and ROI, the beads were modified with 1:1:1 ratio for three capture sequences: *eGFP* : *Pdgfrα*-targeting-exon-7 : *Pdgfrα*-targeting-exon-17 (Supp Figure 10c-d). In addition, eight ROIseq primers were spiked in during library generation, one for *eGFP* detection (ROI^eGFP^) and seven for *Pdgfrα* (ROI^Pα^) to probe splice junctions nearer to the transcript’s 5’-prime end, where usually no information is retrieved in scRNAseq experiments (Supp Figure 10d-e).

We performed the experiment with a 1:1 mixture of sorted eGFP-positive colonic fibroblasts and Epcam-positive epithelial cells (Supp Figure 11a-b), using either unmodified beads (unmod) or RoCK and ROI (rockroi). To simplify the comparison of the WTAs, we combined and sequenced the WTA profiles of the unmod and rockroi libraries in a full cartridge (unimodal condition, WTA and WTA^ROI^ libraries). In a second cartridge, we sequenced the WTA and TSO libraries of the rockroi condition (multimodal condition, WTA^ROI^ and TSO^ROI^). This also removed the effect of the custom sequencing primer, which was only added in the cartridge with the multimodal condition.

As in previous experiments, the WTA sensitivity for the unmod and rockroi samples looked similar in terms of number of genes, number of UMIs and mitochondrial content (Supp Figure 12a-c). We then manually annotated epithelial and fibroblast clusters using known markers (see Brügger et al, 2020^53^; Supp Figure 12d, Supp Table 2). All cell types detected in unmod were also detected in rockroi (Figure 6b), including rare cell types, such as Tuft and enteroendocrine cells. The detected mitochondrial content (Figure 6c) and genes (Figure 6d) across clusters were similar between the unmodified and rockroi conditions.

The ROI^Pα^ primers added during library generation yielded reads spanning splice junctions in the *Pdgfrα* transcript (Figure 6e-f). As expected, these reads were detected exclusively in fibroblast clusters, demonstrating the specificity of the RoCK and ROI method. Additionally, in most of the ROIseq junctions, the percent of positive cells was higher in crypt top compared to crypt bottom cells, which is consistent with previous findings showing that crypt top fibroblasts have a higher *Pdgfrα* (and *eGFP*) expression^53^. In contrast, reads in the regions targeted by ROIseq primers were completely absent in the unmodified sample (Supp Figure 12e-g) but clearly yielded reads spanning the targeted splice junctions in rockroi (Supp Figure 12g). In addition to *Pdgfrα*, we also detected *eGFP* (Supp Figure 13a-c), again with exclusive expression in fibroblasts.

We next compared the cell types detected via the WTAs of the unmod and rockroi conditions to determine if adding a set of ROIseq primers affected the ability to distinguish distinct subpopulations in an scRNAseq experiment. The cell types detected in the unmod and rockroi samples were highly concordant (Figure 6g, Pearson correlation between 0.94 and 0.97), indicating that the addition of multiple ROIseq primers during library generation does not significantly impact the WTA profiles.

We then shifted our focus to the WTA profiles of the unimodal *versus* multimodal rockroi conditions. Since the same library was sequenced twice, this gives a baseline of technical variation; the two WTA readouts were highly correlated (Pearson correlation 0.987, Supp Figure 13d-e).

To discriminate between *Pdgfrα* long and short transcripts, we looked into the discriminant splicing region between exons 16 and 17 (Supp Figure 10b, d-e). First, our RoCKseq capture in exon 17 is specific to the long transcripts. Second, the junction where discriminant splicing occurs is also targeted by the roi_16 primer; reads spanning this exon junction are specific to the *Pdgfrα* long transcripts. The short isoforms on the other hand can be detected by reads mapping to the 3’ UTR of the short *Pdgfrα* isoform, which are present in both rockroi and unmodified samples in crypt bottom and top cells (Supp Figure 13f-g).

Taken together, RoCK and ROI is able to direct reads to specific regions of interest such as splice junctions. By capturing *Pdgfrα* close to a junction of interest and adding a primer spanning this region, RoCK and ROI can also detect and distinguish between splice variants. Furthermore, the WTA profiles detected with RoCK and ROI remain similar and can be used for standard scRNAseq analyses (*e.g.*, cell type annotation).

## Discussion

We present RoCK and ROI, a simple and highly versatile scRNAseq-based method designed to capture specific transcripts (RoCKseq) and to selectively sequence regions of interest (ROIseq). RoCKseq works through the modification of standard barcoded beads, while ROIseq is mediated by addition of primers during library generation. Several quality checks help to assess the performance of RoCK and ROI prior to and throughout the experiment. We also provide a tailored data analysis workflow to systematically assimilate the targeted (TSO) and untargeted (WTA) reads.

RoCK and ROI offers a rapid and reliable bead modification protocol (about two hours) that is titratable and can be multiplexed. Furthermore, the bead modification is stable over months, allowing users working with time-sensitive material (for example clinical samples) to perform and validate RoCKseq bead modification prior to knowing the date of the future experiments. Additionally, RoCKseq capture is highly flexible and may be adaptable to other platforms. We have shown that dT oligos, which are used by most beads on scRNAseq platforms, can be modified using the same protocol designed for the TSO oligos (Supp Figure 3c-d). The potential hurdle to adapt RoCKseq bead modification to other platforms is the bead chemistry, which may not be suited to heating or to the buffers used during the modification protocol. Finally, we show that RoCKseq allows for an accurate titration of modification on the beads, which is guaranteed by the exonuclease step.

RoCKseq capture also leads to a shift in the position within the transcript where reverse transcription is initiated. This is an advantage over targeted amplification methods where reverse transcription occurs at the 3’ end. RoCKseq thus offers the unique possibility to reverse transcribe regions even at the 5’ region of long transcripts, or to avoid GC-rich stretches where reverse transcription is often impaired. In combination with ROIseq primers, a defined cDNA product can be generated that is not only suited for PCR during library preparation but also compatible with the downstream sequencing process. In contrast, targeted amplification approaches are performed after reverse transcription and thus suffer from the biases that already occur during mRNA capture^12^. Moreover, RoCK and ROI is multimodal, since the WTA is profiled alongside the targeted readouts deriving from the TSO data.

Interestingly, we observe that the information retrieved in RoCK and ROI is not exclusively derived from the targeted mRNAs. We show that both the RoCKseq capture and the ROIseq primers target other transcripts beyond the desired ones. This is due to the experimental conditions used during mRNA capture and during library preparation that remain as provided and suggested by the standard protocol (*i.e.*, two minutes at room temperature using ice-cold buffer). The standard parameters are adapted to suit the large diversity of transcripts differing for instance in length, GC-content, or sequence complexity and are optimized to capture polyadenylated mRNAs. However, as a consequence, these relaxed mRNA capture conditions eventually lead to the detection of non-polyadenylated transcripts via dT-capture, which may sum up to 20% of the detected transcripts^54,55^. One possible explanation is that this may occur through the binding to internal polyA sequences present in transcripts. Given that the (modified or unmodified) TSO oligos have higher melting temperatures (T_m_: 63.5°C) than the dT sequence (37.5 °C for a stretch of 25 dTs as present on BD Rhapsody beads), it is no surprise that a variety of non-targeted transcripts is recovered also in the TSO library. Hence, on-target enrichment expectations have to be taken into consideration during sequencing planning. The low percentage of 0.5% of specifically captured transcripts present in the TSO readouts can be explained by the low number of molecules that can be targeted in the total pool of RNA molecules present in a cell: the RNAScope experiment shows that on average 142 targetable *eGFP* mRNAs are present in our clonal L-cells corresponding to 0.014% of the total pool of 10^5^ to 10^6^ mRNAs estimated to be present in mammalian cells^56^. In addition, less than 10% of the RNA molecules present in a cell are polyadenylated^57,58^, and the targetable molecules in the experiment are thus between 14 and 140 per millions of RNAs per cell. At the molecular level, we believe (for both dT and even more for TSO-based capture) that the partial binding of a subset of nucleotides at the 3’ end of the capture oligo (which are about 10^7^ / bead) is sufficient to trap transcripts other than the targeted ones and that a perfect match of a few bases at the 3’ end can initiate reverse transcription. This is in line with the standard use of random hexamers for initiating the reverse transcription^59,60^. An improvement of the on-target RoCKseq capture (as well as ROIseq-based second strand synthesis) would require a change in the conditions of the standard capture and library preparation to increase the binding specificity of RoCK and ROI oligos to the targets. As a direct negative consequence, such changes (*e.g.*, elevated temperatures or adapted buffers) will likely impair the dT-based capture of mRNAs that occur simultaneously on the cell lysate. While only a small fraction of reads from the TSO library are on target (Supp Figure 9a-l), it is still sufficient to obtain the information for the targeted transcripts in many cells.

RoCK and ROI is suited for applications in which users are interested in reading one or multiple specific transcripts. We have shown that RoCKseq capture can be multiplexed, leading to the possibility of multiple transcripts being captured at the same time as in the experiment on murine colonic cells. Multiplexing of RoCKseq capture, on the other hand, leads to a decreased detection rate for each individual transcript as a lower modification rate is achieved.

RoCK and ROI is suitable for a multitude of applications. Any change on the DNA level that is transcribed into RNA, polyadenylated or not, can be investigated using the RoCK and ROI workflow. The list of genetic features that can be analyzed is diverse and ranges from genetically engineered genes, inducible ectopic gene activation, transgenes, Cre-based recombination, naturally occurring sequence variations, or CRISPR screens.

In summary, we believe that the RoCK and ROI workflow is a widely applicable and important addition to the wealth of existing single cell transcriptome sequencing tools. It will help to explore and better understand complex biological systems in health and disease as it enables the detection of specific transcripts or sequence variations in the context of transcriptional phenotypes at the single cell level.

## Methods

### Design of capture sequences and fluorescent oligos

Detailed information on the design of ROIseq primers is available on protocols.io (dx.doi.org/10.17504/protocols.io.rm7vzjyb5lx1/v1).

A list of primers used for the scRNAseq experiments can be found in Supp Table 3.

The sequence of the splint for the modification of TSO oligos was as follows: 5’ -24 nt coding sequence followed by a constant sequence-3’: 5’-NNNNNNNNNNNNNNNNNNNNNNNNCATACCTACTACGCATA-3’ where the CATACCTACTACGCATA is the reverse complement of the TSO sequence. The sequence of the splint acts as a template for capture synthesis. For the modification of dT oligos on beads, the reverse complement of the TSO sequences was substituted by a polyA stretch of 18 nts: 5’-NNNNNNNNNNNNNNNNNNNNNNNNAAAAAAAAAAAAAAAAAA-3’.

The polyA protective oligo used on the barcoded beads was 18 nucleotides in length: 5’-AAAAAAAAAAAAAAAAAA-3’.

The oligos were ordered in 0.2 µmol scale, HPLC grade, with 5’ phosphorylation. Before use, the oligos were resuspended in ddH_2_O to generate a 100 µM stock solution.

Fluorescent oligos were designed by taking the first 20 nucleotides from the 5’ end of the splint. The fluorescent oligos were ordered in HPLC grade and in 0.2 µmol scale with a 5’ Atto647N modification and diluted in ddH2O to generate a 100 µM stock solution.

### Protocol for polymerase-based bead modification for BD Rhapsody beads

A step-by-step protocol is available on protocols.io (dx.doi.org/10.17504/protocols.io.rm7vzjyb5lx1/v1). A general workflow is described in this section. Briefly, in a first step of the bead modification, BD Rhapsody beads (“Enhanced Cell Capture Beads V2”, Part Number 700034960, BD Rhapsody^TM^ Enhanced Cartridge Reagent Kit, BD 664887) were incubated with a splint, protective polyA oligo and T4 DNA polymerase mix (Thermo scientific EP0061) without the enzyme for 5 minutes at 37° C with shaking at 300 rpm. The T4 polymerase enzyme was then added and the mix was incubated for 10 minutes at room temperature with rotation. This was followed by inactivation of the T4 polymerase by incubating the mix for 10 minutes at 75°C. The single-strandedness of the DNA oligos on the beads was restored by incubating the beads with a lambda exonuclease mix (NEB M0262L) for 30 minutes at 37° C, followed by inactivation of the enzyme by incubation for 10 minutes at 75° C. The bead modification protocol was performed on a full vial of BD Rhapsody beads (2 mL) or a small subset of beads (20 µL) to test the splint prior to the scRNAseq experiment.

### Protocol for fluorescent assay to quantify bead modification efficacy by FACS analysis

A step-by-step protocol is available on protocols.io (dx.doi.org/10.17504/protocols.io.rm7vzjyb5lx1/v1). A general workflow is described in this section. To test RoCKseq bead modification, barcoded beads were incubated with multiple fluorescent oligos acting either as positive and negative controls or specific for the modification. RoCKseq modified beads and unmodified beads used for controls were incubated with fluorescent oligos for 30 minutes at 46° C in BD Rhapsody lysis buffer (part number 650000064, BD Rhapsody^TM^ Enhanced Cartridge Reagent Kit, BD 664887) with 1 M DTT (part number 650000063,BD Rhapsody^TM^ Enhanced Cartridge Reagent Kit, BD 664887).

Recommended conditions for the fluorescent assay are as follows:

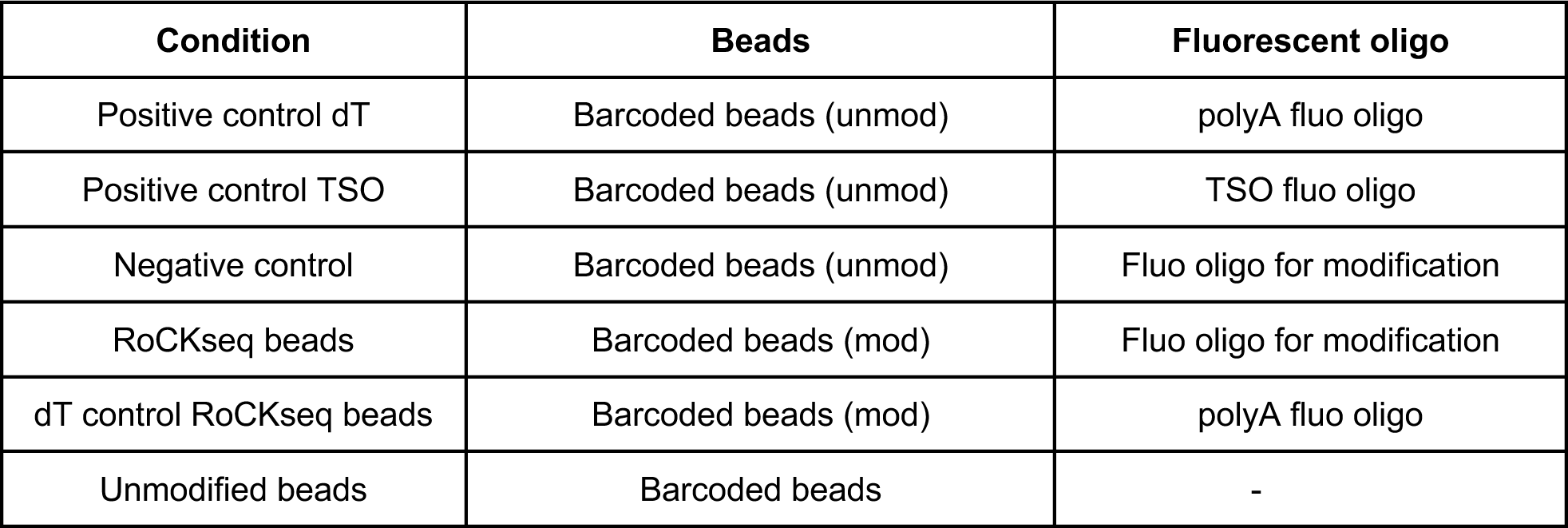

| Condition | Beads | Fluorescent oligo |
| --- | --- | --- |
| Positive control dT | Barcoded beads (unmod) | polyA fluo oligo |
| Positive control TSO | Barcoded beads (unmod) | TSO fluo oligo |
| Negative control | Barcoded beads (unmod) | Fluo oligo for modification |
| RoCKseq beads | Barcoded beads (mod) | Fluo oligo for modification |
| dT control RoCKseq beads | Barcoded beads (mod) | polyA fluo oligo |
| Unmodified beads | Barcoded beads | - |

### Analysis of fluorescent signal from barcoded beads

The signal from barcoded beads after the fluorescent assay was measured at the Cytometry Facility at the University of Zürich using a FACS Canto II 2L with HTS (BD Biosciences, Switzerland). The signal from the Atto647N molecules was measured using the APC-A channel. Gating for beads was performed on the FSC-A versus SSC-A scatterplot and 1000 beads per condition were measured. The .fcs files obtained from the analyser were imported into R (version 4.3.1) and plots were made primarily using the flowCore (version 2.14.0), flowViz (version 1.66.0), ggcyto (version 1.30.0) and ggplot2 (version 3.4.4) packages.

### ROIseq primer design

Detailed information on the design of ROIseq primers is available on protocols.io (dx.doi.org/10.17504/protocols.io.rm7vzjyb5lx1/v1). ROIseq primers were designed directly 5’ (max. 10bp upstream) to the region of interest (ROI). The length of the primers we used is 12 nucleotides. Since 12 nucleotides will be included in the cDNA sequencing read (HTS), the ROIseq primer was designed in close proximity to the ROI. The ROIseq primer has the following structure: 5’-TCAGACGTGTGCTCTTCCGATCTNNNNNNNNNNNN-3’; the N12 sequence of the ROIseq primer is identical to the coding strand. The primers were ordered from Microsynth in HPLC grade and at 0.2 µmol scale and resuspended in DNA Suspension buffer from Teknova (T0221).

### Library generation for RoCK and ROI

A step-by-step protocol is available on protocols.io (dx.doi.org/10.17504/protocols.io.rm7vzjyb5lx1/v1). A general workflow is described in this section, mRNA capture and cDNA synthesis were performed following the manufacturer’s instructions (Doc ID: 210966) using the following kits: BD Rhapsody^TM^ Enhanced Cartridge Reagent Kit: BD 664887; BD Rhapsody^TM^ Cartridge Kit: BD 633733; BD Rhapsody^TM^ cDNA Kit: BD 633773; BD Rhapsody^TM^ WTA Amplification Kit: BD 633801. To account for the bead loss during modification, RoCKseq beads were resuspended in 680 µl Sample Buffer (Cat. No. 650000062, BD Rhapsody^TM^ Enhanced Cartridge Reagent Kit, BD 664887) instead of 750 µl.

The RoCK and ROI libraries were generated following the manufacturer’s recommendations (Doc ID: 23-21711-00) with the following changes:

1. <u>Random priming and extension:</u> ROIseq primers were added after beads were resuspended in Random Priming Mix. If a single ROIseq primer was added, 1 µl of the 100 µM primer was diluted 1:10 in ddH2O and 4 µl of the diluted mix was added. If multiple ROIseq primers were used, 1 µl of each ROIseq primer (100 µM) was mixed, ddH_2_O was added up to 10 µl and 4 µl of the diluted mix was added.
2. <u>RPE PCR</u>: during RPE PCR, 1 µl of 100 µM T primer was added to each sample of RPE PCR Mix combined to Purified RPE product.
3. <u>Indexing PCR:</u> for indexing of RoCKseq libraries, a separate PCR was performed substituting 5 µl of the Library Forward Primer with 5 µl of 100µM of a custom indexing primer. The same primary library and reverse primers were used as recommended by the manufacturer.

If no ROIseq was being performed, points 2-3 were followed. A list of primers used for the scRNAseq experiments can be found under Supp Table 3. The T primer and Indexing primer were resuspended in DNA Suspension buffer from Teknova (T0221).

### Sequencing

Libraries were indexed using the BD Rhapsody Library Reverse primers as described by the manufacturer combined either with the BD Rhapsody Library Forward primer for the WTA-based information or the RoCKseq Indexing primer (see section Library Generation for RoCK and ROI). RoCKseq and dT-based libraries of a given sample were indexed with the same 8 bp index sequence and pooled in a 1:1 concentration. For sequencing of pooled libraries including at least one RoCKseq modified sample (with or without ROIseq primers), a custom R1 primer was spiked in for the sequencing. Sequencing was performed at the Functional Genomics Centre Zurich (FGCZ) using a Novaseq 6000 and a full SP 200 flow cell for each experiment. The length of R1 was 60 bp and the length of R2 was 62 bp. A 3% PhiX spike-in was used.

### Generation of stable cell lines

The FUGW plasmid (Addgene #14883) was used for the generation of the L-cells expressing eGFP. For the generation of the HEK293 cells expressing tdTomato, the eGFP ORF in the FUGW plasmid was excised using the EcoRI and BamHI sites and substituted with the tdTomato sequence from the pCSCMV: tdTomato vector which was excised with the same restriction enzymes. The fluorescent cells were generated by lentiviral transduction. Lentiviruses were generated following the cultured Lipofectamine 3000 protocol supplied by the manufacturer. HEK293T cells were in a T75 flask using 16 mL of packaging medium which was generated by mixing 47.5 mL Optimem reduced serum, 2.5 mL FBS, 100 µl sodium pyruvate and 500 µl Glutamax. On day 1, tube A was prepared by mixing 2 mL of Optimem with 55 µl of lipofectamine 3000. Tube B was prepared by mixing 2 mL Optimem, 17.8 μg of lentiviral packaging plasmid mix (4.8 μg pVSV-G, 9.6 μg pMDL, 3.4 μg pRev), 6 µg of the GFP or tdTomato diluted plasmid and 47 µl of the P3000 reagent. Tube A and tube B were mixed and incubated at room temperature for 20 minutes. 4 mL of medium was removed from the flask and substituted with the 4 mL of mix A and B. The cells were incubated for 6 hours, after which the medium was removed and substituted with 16 mL packaging medium. On day 2, 24 hours post transfection the volume of supernatant was collected and stored at 4 °C. The next day, 52 hours post transfection the medium was collected and stored in the same Falcon tube as the day before. The medium was spun down 2000 rpm for 3 minutes, after which it was filtered through a 45 µm filter into a new tube. The volume was then transferred to an Amicon tube and centrifuged 3000 g for 10 minutes until the volume reached 500 µL on the Amicon tube. The liquid was then stored at -80°C.

For lentivirus transduction, 300’000 HEK or L-cells were seeded onto a 6 well plate 12 hours prior to transduction with 100 uµl of concentrated viral supernatant in standard cell culture medium supplemented with 20µg/mL polybrene (Sigma). The cells were then passaged 3 times.

To generate clonal cell lines, single cells were sorted into single wells of a 96 well plate. For the sorting of the cells, the cells were first of all dissociated with Trypsin-EDTA as described above. The cells were washed once with PBS and spun down at 290x g for 5 minutes. A Zombie Violet viability staining was performed by resuspending the cells in 1 mL PBS and adding 2 µl of Zombie Violet (1:1000 dilution, Biolegend). The cells were then kept for 10 minutes in the dark, after which 9 mL of medium was added to quench the reaction. The cells were then spun down at 290g for 5 minutes, resuspended in 500 mL of medium and filtered through a Falcon 5mL Round Bottom Polystyrene Test Tube with Cell Strainer Snap Cap (352235, Corning). Single cells were sorted in single wells of a 96 well plate at the Cytometry Facility at the University of Zürich using a BD S6 5L cell sorter (BD Biosciences, Switzerland). The cell lines were then expanded and cultured as described above.

### Cell culture

L-cells cells were cultured in Dulbecco’s Modified Eagle Medium 1X (41966-029) with 10% FBS (GIBCO, 10270-106) and 1% PIS in a 10 cm dish and maintained in an incubator at 37°C and 5% CO_2_. Cells were split at 80% confluence. To dissociate the cells, the medium was removed and 2 mL of Trypsin-EDTA (0.5%, no phenol red) were added to the dish, followed by 5 minutes in the incubator. The trypsin was inactivated by adding 8 mL of medium. To remove trypsin, cells were centrifuged for 5 minutes at 290 g, the supernatant was removed and the pellet was resuspended in 10 mL of medium and plated depending on the wanted confluency. The total volume of the dish was 10 mL.

HEK293 were also cultured in Dulbecco’s Modified Eagle Medium 1X (41966-029) with 10% FBS (GIBCO, 10270-106) and 1% PIS in a 10 cm dish and maintained in an incubator at 37°C and 5% CO_2_. Cells were split at 80% confluence. To dissociate the cells, the medium was removed and 2 mL of Trypsin-EDTA were added to the dish. The Trypsin-EDTA was immediately removed and the dish was placed in the incubator at 37°C and 5% CO2 for 1 minute. 10 mL of medium was then added to the dish and the cell mixture was plated depending on the wanted confluency.

### Preparation of single cell suspension from cell lines for scRNAseq experiments

Single cell solutions were prepared following the manufacturers recommendations. After dissociation and spinning down, the medium was removed and the cells were resuspended in 1 mL of Sample Buffer. The cells were filtered through a Round Bottom Polystyrene Test Tube with Cell Strainer Snap Cap (352235, Corning) and counted with a Neubauer chamber. The volume of cell solution to use was calculated using the following formula per manufacturers recommendations: (#cells in experiment x #samples x 1.36) / counted # of cells per µl. The calculated volume was then diluted to 650 µl of sample buffer before loading on the BD Rhapsody Express machine. For the mixing experiments, the same procedure as above was used and the same number of cells were mixed in a 1:1 ratio for a final volume of 1.3 mL.

### Mice and ethics statement

We affirm to have complied with all relevant ethical regulations for animal testing and research as follows. All animal based experimental procedures at the University of Zurich were performed in accordance with Swiss Federal regulations and approved by the Cantonal Veterinary Office (license ZH045/2019). Mice from the *Pdgfrα^H2BeGFP^* strain^52^ were purchased from Jackson Laboratories, United States of America (strain number 007669).

Mice in the *Pdgfrα* scRNAseq experiment were three males aged 2 months and 12 days (for two mice) and 1 month and 22 days.

### Sequencing of transgenic *Pdgfr****α*** locus

To gain the sequence information for mapping of reads in the *Pdgfrα^H2BeGFP^* strain, DNA from tail biopsies was PCR amplified using primers outside of the region removed during generation of the mouse strain^52^, corresponding to a 6.5 kb fragment between BamHI-SmaI sites; sequences of primers forward: ACAGAGGCTGCCTCAAAGCTAG, reverse: CCATTGCCCAGATGGGAAGC) and cloned into pGEM-T easy vector (Promega). The insert was Sanger sequenced using M13 forward and reverse primers. Sequencing was performed by Microsynth.

### Colonic single cell isolation and cell sorting

Colonic tissues were obtained from *Pdgfrα^H2BeGFP^* reporter mice^52^. The tissues were flushed with PBS, longitudinally opened and finely minced into 2 mm pieces. Minced tissue fragments were washed with PBS three times. Following the methodology outlined by Brügger et al, 2020^53^, tissue pieces underwent rounds of digestion to separate epithelial and mesenchymal fractions.

For the detachment of the epithelial fraction, the tissue pieces were incubated in Gentle Cell Dissociation Reagent (STEMCELL Technologies, Germany) while gently rocking for 30 minutes at room temperature. The pieces were pipetted up and down for the epithelial fraction to be detached. The epithelial fraction was then filtered through a Falcon 70-μm cell strainer (Corning, Switzerland), washed with plain ADMEM/F12 and incubated for 5 minutes at 37°C in prewarmed TrypLE express (Gibco, Thermofisher, Switzerland). The gentleMACS Octo Dissociator (Miltenyi Biotec, Switzerland) m_intestine program was employed for single-cell dissociation. The obtained epithelial single-cell suspension was then filtered through a Falcon 40-μm cell strainer (Corning) and kept on ice in ADMEM/F12 supplemented with 10% FBS.

For dissociation of the mesenchymal fraction, the remaining tissue pieces (following epithelium detachment) were digested for 1 hour at 37°C under 110 rpm shaking conditions in DMEM supplemented with 2 mg/mL collagenase D (Roche) and 0.4 mg/mL Dispase (Gibco). The mesenchymal fraction was then filtered through a Falcon 70-μm cell strainer (Corning), washed with plain ADMEM/F12, and subsequently filtered through a Falcon 40-μm cell strainer (Corning).

The epithelial and mesenchymal cells were mixed and stained for 30 minutes on ice with anti-CD326(EpCAM)-PE-Cy5 (1:500, eBioscience/Thermofisher, Switzerland) in PBS. Prior to cell sorting, all cells were stained for 5 minutes on ice with DAPI in PBS (1:1000, ThermoFisher, Switzerland). Epithelial and mesenchymal cells labeled with PE-Cy5 and eGFP were sorted separately and subsequently mixed in a 1:1 ratio. Cells were sorted at the Cytometry Facility at the University of Zürich using a FACSAria III cell sorter (gates visible in corresponding figures) (BD Biosciences, Switzerland).

### RNAScope experimental procedure

The localisation of eGFP mRNAs in cells was performed with RNAScope (Advanced Cell Diagnostics, Germany) in a 96 well plate following the manufacturers recommendations (RNAscope Fluorescent Multiplex Assay). The fluorescent Probe - EGFP-O4 - Mycobacterium tuberculosis H37Rv plasmid pTYGi9 complete sequence (Advanced Cell Diagnostics, 538851) was used for all experiments. After DAPI staining, a protein stain was performed using Alexa Fluor™ 488 NHS-Ester (Succinimidylester) (Thermo Scientific, A20000). The wells were first of all washed with PBS, after which the supernatant was aspirated to 30 µl and 160 µl of CASE buffer (609.4 µl freshly thawed NaHCO_3_, 15.63 µl Na_2_CO_3_, 2.5 mL of water) was added to each well and subsequently aspirated to 30 µL 0.5 µL of Alexa Fluor™ 488 NHS-Ester (Succinimidylester) were then added to the remaining CASE buffer and 30 µl of CASE stain were added to each well. The plate was incubated for 5 minutes at room temperature in the dark, followed by 4 washes with PBS.

### RNAScope image acquisition and analysis

Images were acquired using an automated spinning disk microscope Yokogawa CellVoyager 7000 equipped with a 60x water-immersion objective (1.4 NA, pixel size of 0.108 mm), 405/488/647 nm lasers, the corresponding emission filters and sCMOS cameras. 45 *z*-slices with 0.5 mm spacing were acquired per site. Image analysis was conducted with MATLAB (R2021b) and its image processing toolbox. Raw images were corrected for non-homogeneous illumination for each channel by dividing each pixel intensity value by its normalized value obtained from images of the corresponding fluorophores in solution. Cell segmentation was performed using the maximum-projected Succinimidyl ester staining channel and cellpose^61^ using cyto2 model and a cell diameter of 200 pixels. Segmented cells touching image borders, smaller than 10^3^ or larger than 10^5^ pixels were discarded for further analysis. FISH channel was smoothed using a 3D Gaussian filter (s = 1 pixel) and FISH spots with *x* and *y* coordinates overlapping with segmented cells were detected in 3D using intensity thresholding (100 grays level value) followed by watershed segmentation with a minimum size of 9 voxels. Images were processed with ImageJ (Fiji version 2.0.0-rc-69/1.52p). Maximum intensity projections of 45 stacks are shown.

### BD Rhapsody barcode structure

Our data analysis workflow relies on mining the dual oligos present on beads from BD Rhapsody. Namely, whole transcriptome analysis (WTA) oligos profiling the non-targeted transcriptome have a tripartite cell barcode and a 8-nt-long UMI structure as follows: prepend-N{9}-GTGA-N{9}-GACA-N{9}-UMI with a prepend to choose from none, T, GT or TCA. Template-switching oligos (TSO), modified via RoCKseq, are shaped N{9}-AATG-N{9}-CCAC-N{9}-UMI, without a prepend. The fixed parts between cell barcode 9-mers allow targeted (TSO) from untargeted (WTA) data to be distinguished.

### Single-cell data analysis workflow

We have developed a method to automate data processing from raw reads to count tables (and R SingleCellExperiment objects) and descriptive reports listing both on-target TSO (the targeted data) and off-target WTA (whole transcriptome analysis, the untargeted dT-captured mRNAs) readouts. The software stack needed to run the method is provided via system calls (compiling recipes are provided), conda (environment files provided) or via Docker containers. The workflow is written in Snakemake^46^.

To analyze their data, users need to provide their sequencing files in compressed FASTQ format (one file for the cell barcode plus UMI; and another for the cDNA) and a configuration file specifying the experimental characteristics and extra information, including:

- A genome (FASTA) to align the genome to (*i.e.,* hg38, mm10 etc). The genome needs to contain all (on target) captured sequences, so if these do not belong to the standard genome (*i.e.*, GFP, tdTomato), the genome FASTA file needs to be updated to append the extra sequences.
- Gene annotation (GTF) whose features are quantified separately for WTA and TSO. It is expected to contain a whole transcriptome gene annotation (*i.e.,* Gencode, RefSeq etc) as well as an explicit definition of the RoCK and/or ROI targets captured by the TSO. Instructions to build this GTF are included within the software’s documentation.
- A set of cell barcode whitelists following BDRhapsody’s standards (standard BDRhapsody cell barcodes are included within the software)
- Parameters to fine tune CPU and memory usage.

The workflow (depicted in Figure 2a) follows these steps:

1. Index the reference genome with STAR^47^.
2. Subset reads match the WTA cell barcodes and map those to the transcriptome (genome plus GTF) using STARsolo^48^. Detected cell barcodes (cells) are filtered in at two levels: first, by matching to the user-provided cell barcode whitelist; and second, by applying the EmptyDrops^62^ algorithm to discard empty droplets. We report two outputs from this step: the filtered-in cells according to the aforementioned filters; and the unbiased, whole-transcriptome WTA count table.
3. Subset reads matching both the TSO CB structure and the filtered in cell barcodes and map those to the transcriptome. Our reasoning is that the expected TSO transcriptional complexity is undefined and not usable to tell apart cells from empty droplets, so we borrow the filtered-in cells from the EmptyDrops results from the WTA analysis.
4. (optional) Count on-target features in a more lenient way, filtering in multioverlapping and multimapping reads. This run mode is recommended when the captured regions target non unique loci (*i.e.,* repetitive sequences).

Hence, our workflow always reports a WTA count table with as many genes as on-target and off-target gene features in the GTF, and per filtered-in cell barcode. As for the TSO, we offer these run modes:

- *tso off- and ontarget unique*: generates a count table for TSO reads from filtered-in cells; this count table has the same dimensions as the WTA.

- *tso ontarget multi*: creates a count table for TSO reads from filtered-in cells for only on-target features while allowing for multioverlapping and multimapping alignments.

- *all*: produces both ’tso off- and ontarget uniquè and ’tso ontarget multì outputs.

Finally, we generate an R SingleCellExperiment object with the aforementioned count tables and the following structure:

- *wta* assay: raw counts from the WTA analysis.

- (optional) *tso_off_and_ontarget_unique* assay: raw counts from the ’tso off- and ontarget’ or ’all’ run modes.

- (optional) *tso_ontarget_mult*i altExp alternative experiment: raw counts from the ’tso ontarget multì run mode. A complementary altExp built on WTA data, named ’wta_ontarget_multì, quantifies multioverlapping and multimapping reads to the on-target regions in WTA data.

We also provide a simulation runmode to showcase the method, where raw reads (FASTQs), genome and GTF files are generated for three on-target features and one off-target feature across hundreds of cells before running the method.

Our method is available at https://zenodo.org/records/11070201 under the GPLv3 terms.

### Reference genomes and annotations

To process the mouse and human mixing experiments, we generated a combined genome by concatenating GRCm38.p6 (mouse), GRCh38.p13 (human) and *eGFP* (sequence obtained from FUGW Addgene #14883) and *tdTomato* (sequence obtained from pCSCMV: tdTomato Addgene #50530). For gene annotation, we used GENCODE’s M25 (mouse) and v38 basic (human) and custom GTFs for eGFP and tdTomato. The data from the *Pdgfrα* experiment were mapped using the mouse genome GRCm38.p6and GENCODE’s M25 annotation, as well as the sequence for the *H2B-eGFP* construct in the transgenic mouse strain that was determined by sequencing the locus as described above.

For mixing experiments, two regions were distinguished: the coding sequence (CDS) and the full transcripts (tx), the latter of which contains the 5’ and 3’ UTR in addition to the CDS.

GTF annotations are available under the GEO accession GSE266161.

### Analysis of high-throughput sequencing data

#### Software versions

Data analysis was performed using R (version 4.3.2). Data wrangling was mainly performed using dplyr v1.1.4 and reshape2 v1.4.4. Plots were generated with ggplot2 v3.4.4 and ggrastr v1.0.2. Omics downstream analysis were run mainly using the Bioconductor ecosystem^17^: scran v1.30.2, scuttle v1.12.0, scDblFinder v1.16.0, Gviz v1.46.1, GenomicRanges v1.54.1, GenomicAlignments v1.38.2, GenomicFeatures v1.54.3 and edgeR v4.0.16. Alignment statistics were retrieved with Qualimap2^63^ v2.3.

#### Downsampling of single-cell data

When applicable (Supp Figure 5d-f, Supp Figure 6d-f, Supp Figure 12a-b), data were downsampled across samples to the lowest average cell-wise library size using the *downsampleMatrix()* function of the scuttle package. The downsampled data were only used to generate QC plots as well as calculating metrics such as mean number of genes or mitochondrial percent per cell.

#### Single-cell quality control metrics and filtering

Quality control metrics for dT and TSO data such as percent mitochondrial transcripts, total number of genes and total number of transcripts were calculated using *addPerCellQCMetric()* from the scuttle package. Library size factors were calculated using *librarySizeFactors()* from the scuttle package.

Datasets were filtered for total number of UMIs and percent mitochondrial transcripts detected in the dT-based data ((first mixing experiment: unmod total > 3000, unmod_T total > 3700, mitochondrial transcripts for both samples >2% and <28%; second mixing experiment: unmod total >3500, unmod_roi total >3500, rock total >2750, rockroi total >3700, mitochondrial transcripts for both samples >2% and <28%; *Pdgfrα* experiment: for both samples total > 800, mitochondrial transcripts >1% and < 75%). If two species were present in the experiment (such as for the first and second mixing), the filtering was performed based on the sum of percent mitochondrial transcripts for the two species. Additionally, genes having less than three counts detected over all cells were filtered out in dT data.

Doublet removal was performed using scDblFinder^64^ stratified by sample (*e.g.*, rock, rockroi etc). Doublets were filtered out.

#### Species assignment (mouse versus human)

To distinguish between mouse and human cells in the two mixing experiments, we aligned the raw reads against a combined genome including mouse, human and other sequences (see section Reference genomes and annotations). Mouse cells were defined as having more than 50% counts to mouse genes or *eGFP* and *tdTomato* sequences and *vice versa* for human cells. Cells were labeled as “unknown” when having less than 50% of either mouse and human genes and were removed from the dataset for downstream analysis.

#### Generation of coverage plots

Coverage plots for the *eGFP* and *tdTomato* transcripts were generated using UMI-deduplicated BAM files containing both unique and multimapping alignments as generated by the workflow described above. The BAM files were split into mouse versus human cells based on the species assignment described above. Plots were generated using the Gviz package^65^. Ranges for the annotation track were specified using the GenomicRanges and GenomicAlignments.

Coverage plots for ROIseq peaks in other genes were generated based on UMI-deduplicated bigWig files outputted by the workflow described above. Plots were generated using Gviz, as described above. The annotation track was generated by transforming the GTF used for mapping into a TxDb object using GenomicFeatures.

Coverage plots across mitochondrial transcripts were generated based on deduplicated bigWig files outputted by the automated pipeline described above. Plots were generated using Gviz as described above

#### Detection of positive cells (mixing mouse and human experiments)

The percent of positive cells for *eGFP* and *tdTomato* was based on counting UMI-deduplicated, unique or multimapping reads. The number of cells with non-zero counts for the CDS in the appropriate cell type (mouse or human cell line) was divided by the total number of cells after deduplication and filtering.

#### Pseudobulk analysis of WTA signal across beads modifications

For the *Pdgfrα* experiment rockroi versus unmod analysis, to compare the WTA data between conditions, counts deriving from the previously filtered, doublet removed object were first aggregated by calculating the average logcount for each gene over each cluster. Genes with mean logcount across all cells (independent of cluster / condition) higher than 0.1 and variance higher than 0.5. were kept for the analysis. Bead modifications were compared by correlating (Pearson) pseudobulk values pairwise using the built-in *cor()* function from R.

For the two mixing experiments, counts were aggregated using the *aggregateAcrossCells()* function of the scuttle package. Genes with 0 counts across all samples were then removed from the dataset. Logcpm counts were calculated using the *cpm()* function from edgeR (*prior.count=1*). Similarly, conditions were pairwise compared using Pearson correlation with the built-in *cor()* function from R.

For the comparison between the *Pdgfrα* rockroi unimodal and multimodal samples, datasets were subsetted for the same barcodes detected in both samples. Highly variable genes were calculated using the *modelGeneVar()* function (1938 genes with p value < 0.05 ). Counts per million were calculated using the *cpm()* function, after which data were subsetted based on the top 100 most highly expressed of the top 500 variable genes. The Pearson correlation was calculated using the built-in cor() function from R.

#### Calculation of average *eGFP* counts in scRNAseq experiments (RNAScope experiment)

The average *eGFP* counts detected in scRNAseq experiments was calculated based on counting UMI-deduplicated alignments including multimappers. That is, reads aligning to n loci were assigned 1/n counts per locus. These values for the unmod and rockroi conditions were then divided by the sum of the mean RNAScope spots detected per cell for the two eGFP replicates divided by two ((131+118)/2).

#### Gene-body coverage profile plots

Data on the coverage along gene bodies (*e.g.*, from TSS to TES) were generated using rnaqc from Qualimap2^63^. Coverage data were imported into R and plotted using ggplot2.

#### Gene biotypes analysis for TSO data

Gene types detected in TSO data were derived by importing the GTF file used during mapping containing Gencode’s assigned biotypes. The GTF was filtered for genes detected in the WTA from the previously QC-ed, doublet removed object.

#### Sankey diagrams and number of reads and alignments

Data plotted in Sankey diagrams were derived from BAM files generated by the automated pipeline described above. Data on counts (including on-target values) were generated in R. Sankey diagrams were plotted using SankeyMATIC (https://sankeymatic.com/, commit 088a339). The number of reads with canonical WTA and TSO barcode structure were calculated by running a regular expression on FASTQ files and without taking into account the variable regions whitelists. Sankey nodes reporting alignments or counts report our workflow’s outputs, hence taking into account cell barcode whitelists and UMI duplicates. The number of alignments was extracted using bamqc from Qualimap2^63^.

#### Single-cell RNA-seq dimensionality reduction, embedding, and clustering

Dimensionality reduction was performed using WTA-based data after quality control (including doublet removal). First of all the per-gene variance within each condition was modeled using the scran package (*modelGeneVar()* with condition id as block) on log-normalized counts (generated with the *logNormCounts()* from the scuttle package).

Non-mitochondrial genes with biological variance larger than 0.01, p value smaller than 0.01 and mean normalized log-expression per gene were used for dimensionality reduction using the scran package. PCA was calculated with 30 components and used to build UMAP cell embeddings. Cells were clustered using *clusterCells()* from the scran package.

#### Cell annotation (*Pdgfrα* experiment)

Clusters were manually annotated based on known cell markers in Supp Table 2. Cells were first of all split broadly into mesenchymal and epithelial and then clustered independently for annotation. Epithelial and mesenchymal clusters were defined as having mean logcounts per cell higher than 0.35 over all defined epithelial or mesenchymal markers respectively. Logcounts were calculated using the *logNormCounts()* function of the scuttle package. As one epithelial cluster had markers for both enteroendocrine and Tuft cells, the clustering was rerun on the subset of cells to distinguish the two cell types. Cells that were not classified as epithelial or mesenchymal were removed from the dataset.

#### Junction analysis for *Pdgfrα*

To detect reads spanning splice junctions, BAM files for WTA and TSO datasets were split by cell barcode and cell type (crypt top, crypt bottom and epithelial) and counted with featureCounts^66^ specifying the *-J* (junction) flag and using fraction counts for multimappers (*--fraction*). Only canonical (annotated) splice junctions were kept into consideration. Only QC-filtered and doublet removed cell barcodes were included into the analysis.

The coverage, sashimi and alignment tracks for the roi_16 region were generated using Gviz. Only splice junctions with at least one UMI were filtered in.

We refer to GENCODE M25 ENSMUST00000202681.3 and ENSMUST00000201711.3 as short *Pdgfrα* transcripts; and to ENSMUST00000000476.14 and ENSMUST00000168162.4 as long transcripts.

## Data availability

Raw and processed data are available at GEO accession GSE266161.

## Code availability

Source code to analyze data from our method are available at https://zenodo.org/records/11070201 under the GPLv3 terms; and the code used to generate the figures and tables in this manuscript are available at https://zenodo.org/records/11124929 with MIT license.

## Acknowledgments

We thank Vadir López-Salmerón, Cynthia Sakofsky, Hye-Won Song, Jannes Ulbrich, Margaret Nakamoto from Becton Dickinson (BD) for their advice and technical support. We thank Catharine Fournier Aquino, Hubert Rehrauer, Andreia Cabral de Guevea, Hai Bui, Joel Wirz at the Functional Genomics Center Zurich (FGCZ), Mario Wickert and Tatiane Gorski at the Cytometry Facility UZH, as well as Costanza Borrelli, Nidhi Agrawal, Jamie Little, Barbara Hochstrasser, Reto Gerber for their technical support. We also thank George Hausmann for manuscript reading. We thank George Hausmann, Achim Weber, Pierre-Luc Germain as well as the rest of the Robinson lab as well as the Basler lab for scientific discussions. We thank Fabienne Brutscher and Jamie Little for their support. Additionally, we are grateful for reagents received from BD and the Pelkmans lab. Quentin Szabo was supported by an EMBO postdoctoral fellowship (ALTF number: 170-2021) and a SNSF Swiss Postdoctoral Fellowships (TMPFP3_210503). T.V. was partially supported by The project National Institute for Cancer Research (Programme EXCELES, ID Project No. LX22NPO5102) funded by the European Union (Next Generation EU). This work was supported by the Swiss National Science Foundation (SNF), grant numbers 192475 (K. Basler) and 310030_204869 (M. Robinson), and a grant from the Julius Klaus-Stiftung to E. Brunner.

## Author contributions

E. B., K. B., and R. Z. conceived the study; E. B., K. B., I. M. and M. D. R. supervised the study; K. B. and E. B. were responsible for funding acquisition; E. B., G. M., I. M., M. D. R., K. B., R. Z. contributed to experimental design; G. M. developed the wet-lab protocol, performed wet-lab experiments and downstream data analysis; I. M. developed the data analysis pipeline and performed downstream data analysis; M. D. R. performed downstream data analysis; M. D. B. performed the RNAScope experiment; H. F. performed the isolation of murine colonic cells for the scRNAseq experiment; J. M. contributed to protocol development and validation; Q. Z. performed the imaging and computational analysis of the RNAScope experiment; F. K. contributed to protocol validation; K. H. contributed to initial wet-lab protocol validation experiments; T. V. was responsible for mouse crosses, genotyping and licenses; G. M., E. B., I. M. and M. D. R. wrote the manuscript and all co-authors commented and edited it.

## Glossary

AP: Allophycocyanin
BAM: Binary Alignment Map
CB: Cell Barcode
CDS: Coding Sequence
Corr: Correlation
dNTPs: Deoxyribonucleotide triphosphate mix
FSC: Forward Scatter
GTF: Gene transfer format
HTS: High throughput sequencing
mod: modified
mt: mitochondrial
neg: negative
oligos: oligonucleotides
QC: Quality Control
RoCKseq: Robust Capture of Key transcripts
ROI: Region Of Interest
ROIseq: Region Of Interest method
scRNAseq: single-cell RNA sequencing
SSC: Side Scatter
TES: Transcription End Site
TSO: Template Switching Oligo
TSS: Transcription Start Site
tx: transcript
U primer: Universal primer
UMAP: Uniform Manifold Approximation and Projection
UMI: Unique Molecular Identifier
unmod: unmodified
UTR: Untranslated Region
WTA: Whole Transcriptome Analysis

---

**Supp Figure 1:**
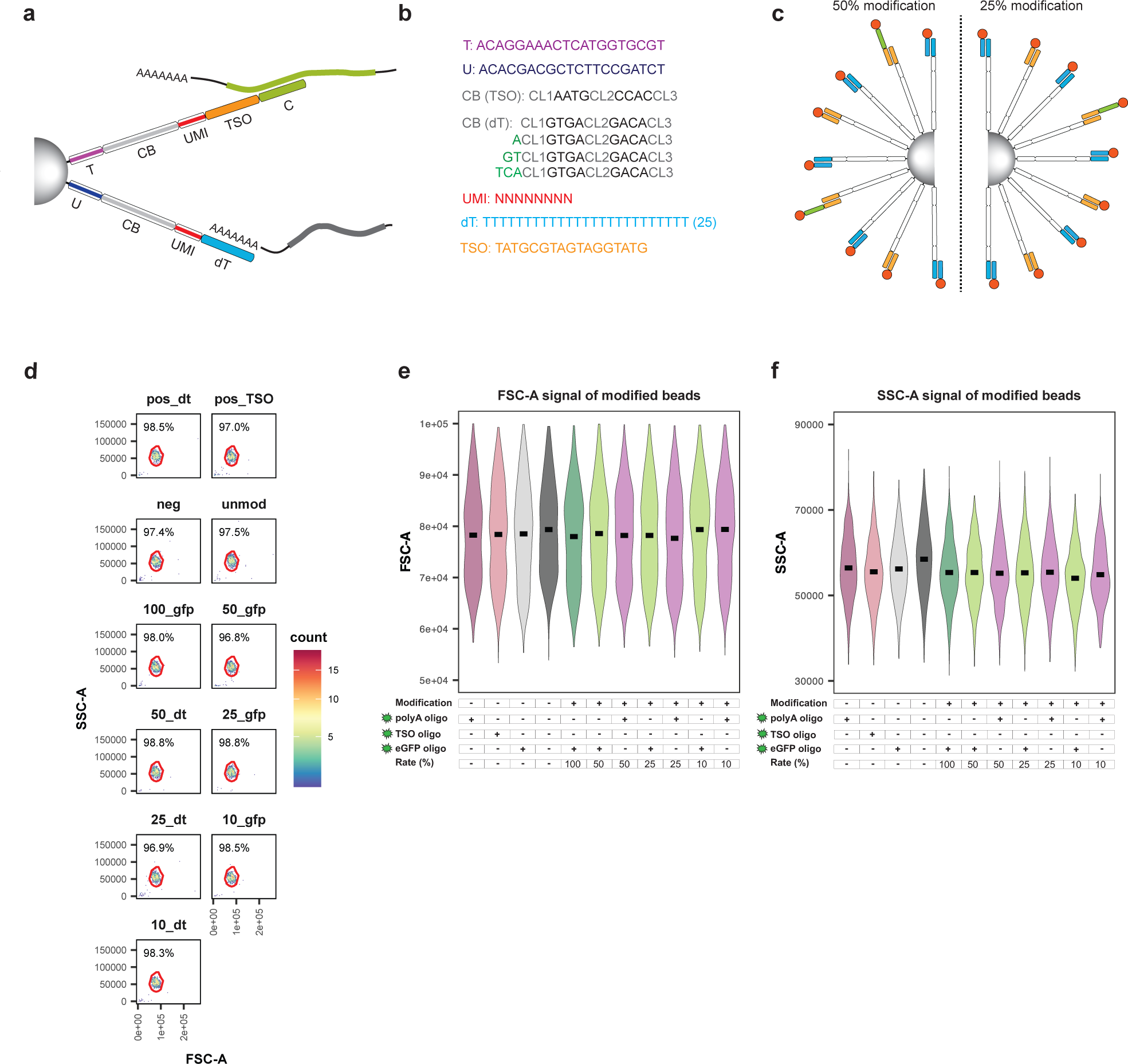
Sequence information on BD Rhapsody barcoded beads and size of RoCKseq modified beads. **a,** RoCKseq BD Rhapsody beads. T: T primer, U: universal primer, CB: cell barcode, UMI: unique molecular identifier, TSO: template switching oligo, C: capture sequence added to the beads. **b,** Bead elements’ sequence: T and U primers; CB (TSO): cell barcodes on TSO oligos; CB (dT): cell barcodes on WTA oligos; UMI; dT (WTA oligos); TSO (TSO oligos). **c,** Titration of RoCKseq modification; left 50% modification, right: 25% modification. The TSO titration oligo is also 5’ phosphorylated (red circle) as it requires removal by the lambda exonuclease. **d-f,** Size of barcoded beads after titration of RoCKseq modification**. For panels (e-f)**: The Y-axis has a biexponential transformation.

**Supp Figure 2:**
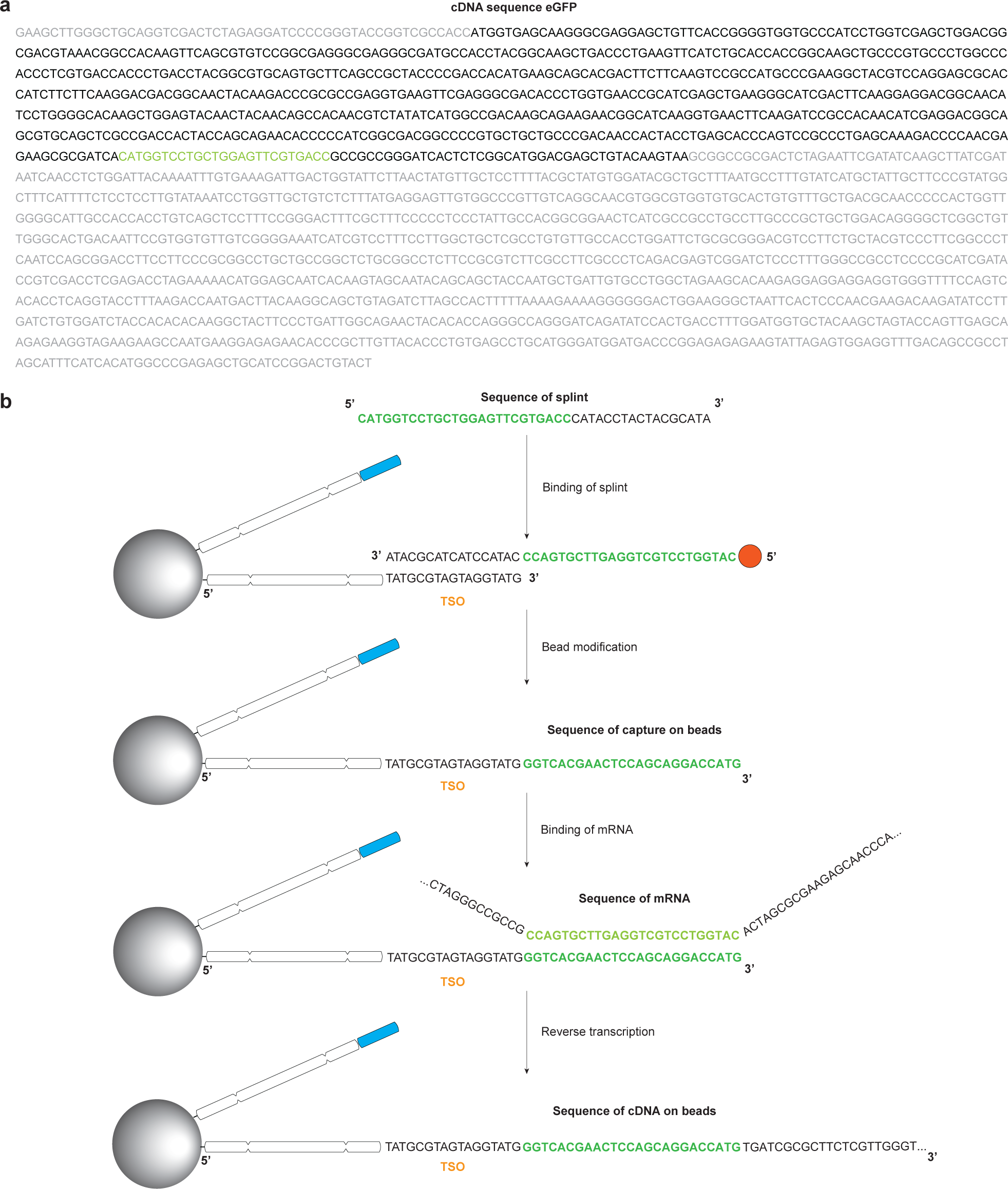
Design of splints for RoCKseq bead modification. **a,** *eGFP* cDNA sequence. Grey: UTRs, black: *eGFP* CDS, green: capture (RoCKseq) sequence. **b,** bead modification process to capture *eGFP*: splint binding, RoCKseq modification and target capture and reverse transcription. The splint contains three elements: a region complementary to the TSO sequence on the beads, the reverse complement of the capture sequence and a 5’ phosphate group (red circle).

**Supp Figure 3:**
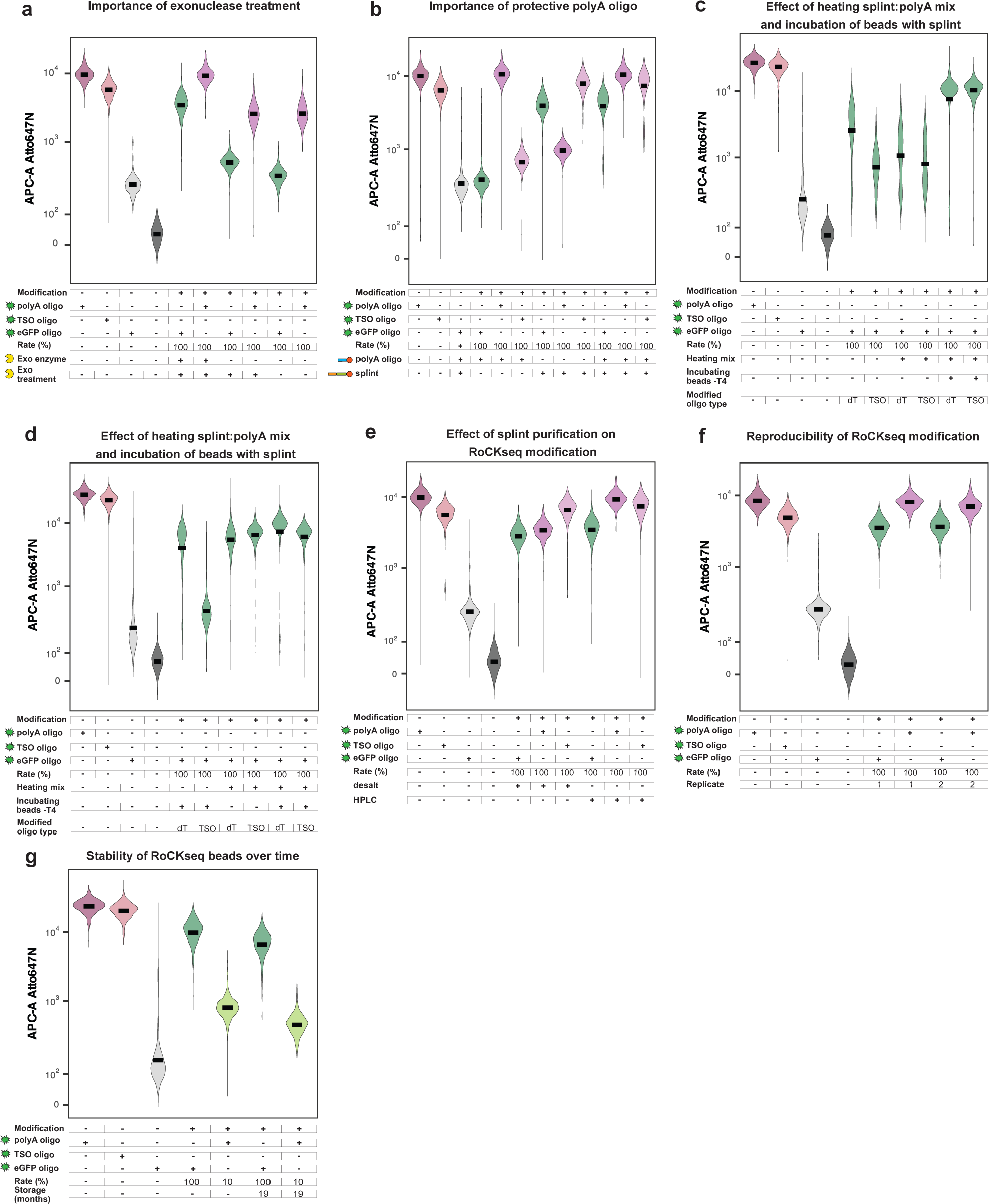
Validation of RoCKseq bead modification. **a-g,** FACS quantification of RoCKseq bead modification. Effect of exonuclease treatment during RoCKseq bead modification **(a)**. Target: *eGFP* CDS. Exo enzyme: addition of exonuclease enzyme to reaction; Exo treatment: exonuclease step (including buffer and water, no enzyme). Effect of T4 polymerase 3’ → 5’ exonuclease activity on barcoded bead oligos **(b)**. Target: eGFP. polyA oligo: only protective polyA oligo used for modification (omission of splint). splint: only splint used for modification (omission protective polyA oligo). Effect of heating of the splint/ polyA mix and incubation of beads with splint before addition of T4 polymerase enzyme **(c)**. Target: *eGFP* CDS. Heating mix: splint/ polyA mix was heated to 75°C for 5 minutes; Incubating beads -T4: beads were incubated with the splint at 37°C for 5 minutes before addition of the T4 polymerase. Modified oligo type: modification of dT or TSO oligos on BD Rhapsody beads. Effect of incubation of beads and splint before addition of T4 polymerase enzyme with or without heating of splint/ polyA mix **(d)**. Conditions as in **(c)**. Effect of purification level of splint and protective oligo on RoCKseq modification **(e)**. Target: *eGFP* CDS. desalt: RoCKseq bead modification with splint in desalted purification; HPLC: RoCKseq bead modification with splint with HPLC purification. Reproducibility of RoCKseq modification **(f)**. Target: *eGFP* CDS. Replicate: technical replicates of RoCKseq bead modification. Storage of RoCKseq beads **(g).** Target: *eGFP* CDS.

**Supp Figure 4:**
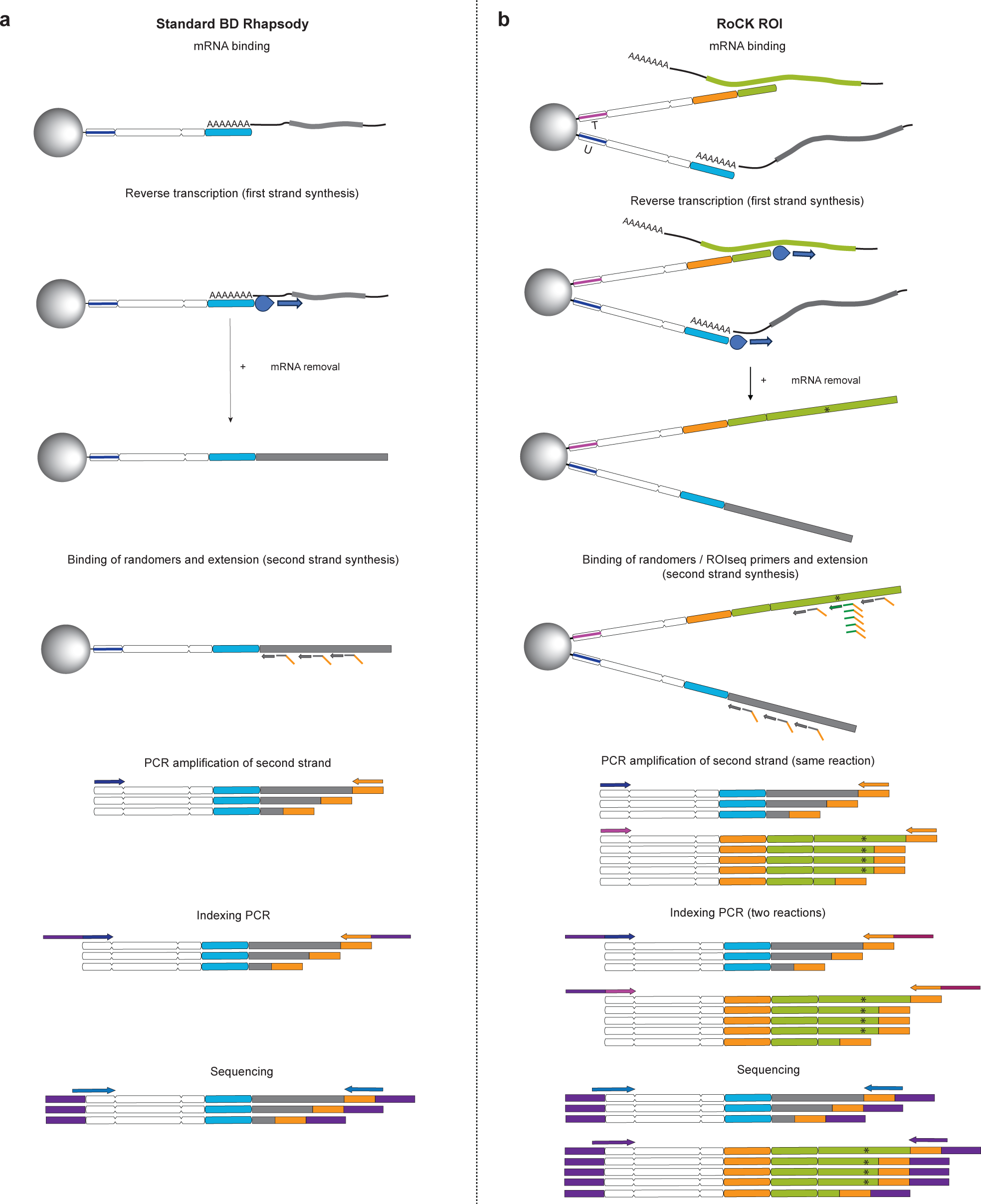
Comparison of RoCK and ROI and standard BD Rhapsody library generation. **a,** Standard BD Rhapsody library generation. **b,** RoCK and ROI library generation using RoCKseq beads.

**Supp Figure 5:**
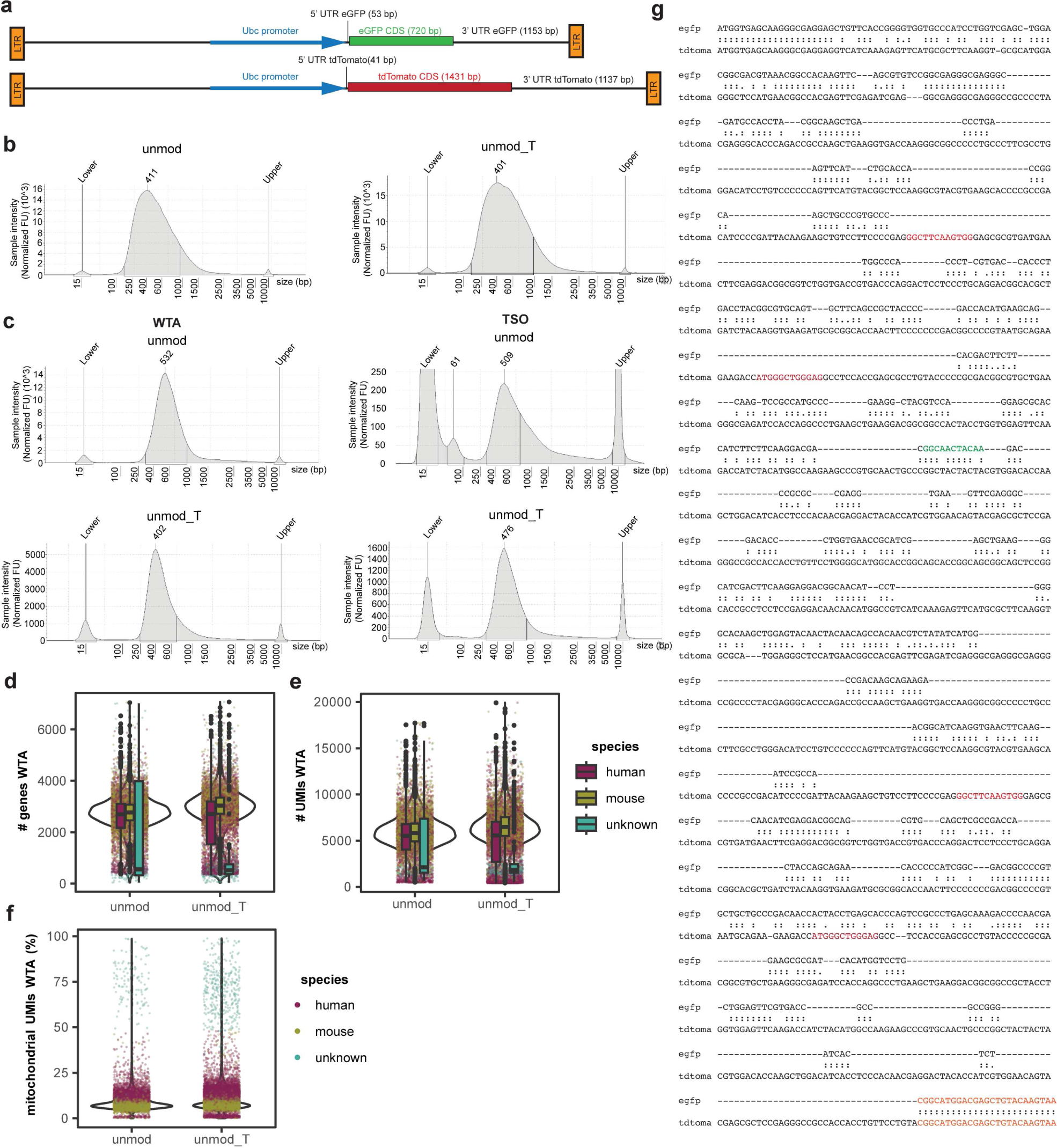
Structure of transgenic construct in cell lines, quality control metrics for scRNAseq experiment to test addition of T primer and position of RoCK and ROI primers in constructs. **a,** *eGFP* and *tdTomato* construct structure. 3’ and 5’ UTRs are identical (differences in number of bases are due to cloning). **b-c,** Library size distribution for unmod and unmod_T samples before **(b)** and after **(c)** indexing. **d-f,** Number of genes **(d)**, UMIs **(e)** and mitochondrial content **(f)** detected in WTA data on downsampled count tables. Figure legend in **(e)** applies to **(d)** and **(e)**. **g,** LALIGN local alignment of *eGFP* and *tdTomato* sequences showing the sequence similarities of the two CDSs. Orange: capture sequence used for the mixing experiments. Red: ROIseq primers for tdTomato. As *tdTomato* is a perfect repeat the ROIseq primers will bind twice. Green: ROIseq primer for *eGFP*.

**Supp Figure 6:**
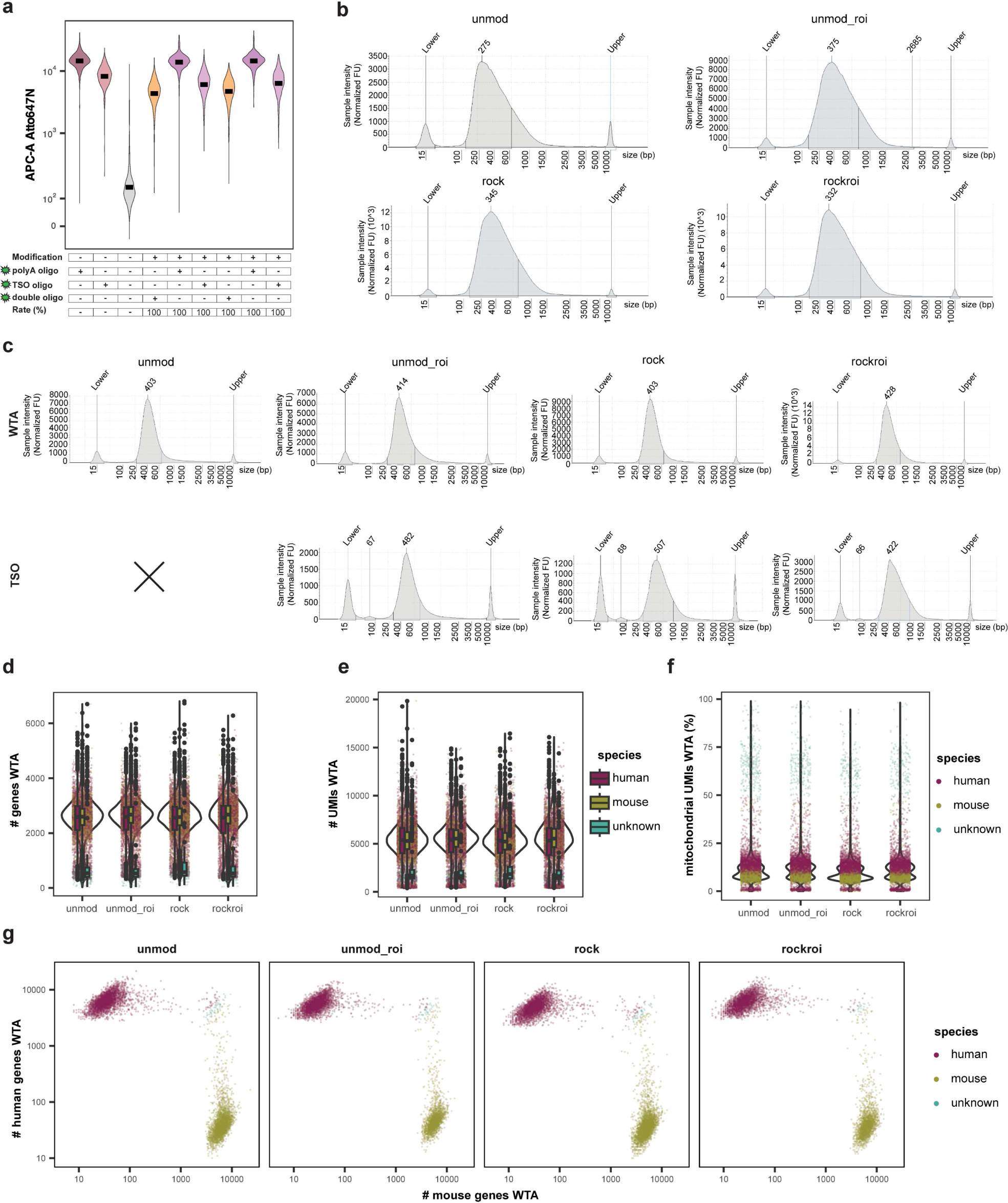
Quality control metrics on WTA data for scRNAseq experiment using mix of cell lines to test RoCK and ROI performance. **a,** FACS signal from modification of beads for scRNAseq experiment. Y-axis: Atto647N fluorescent signal. The Y-axis has a biexponential transformation. **b-c,** Library sizes for unmod, unmod_roi, rock and rockroi samples before **(b)** and after **(c)** indexing. **d-f,** Number of genes **(d)**, UMIs **(e)** and mitochondrial content **(f)** detected in downsampled WTA data. Figure legend in **(e)** applies to **(d)** and **(e)**. **g**, Barnyard plot of species assignment using WTA data per condition.

**Supp Figure 7:**
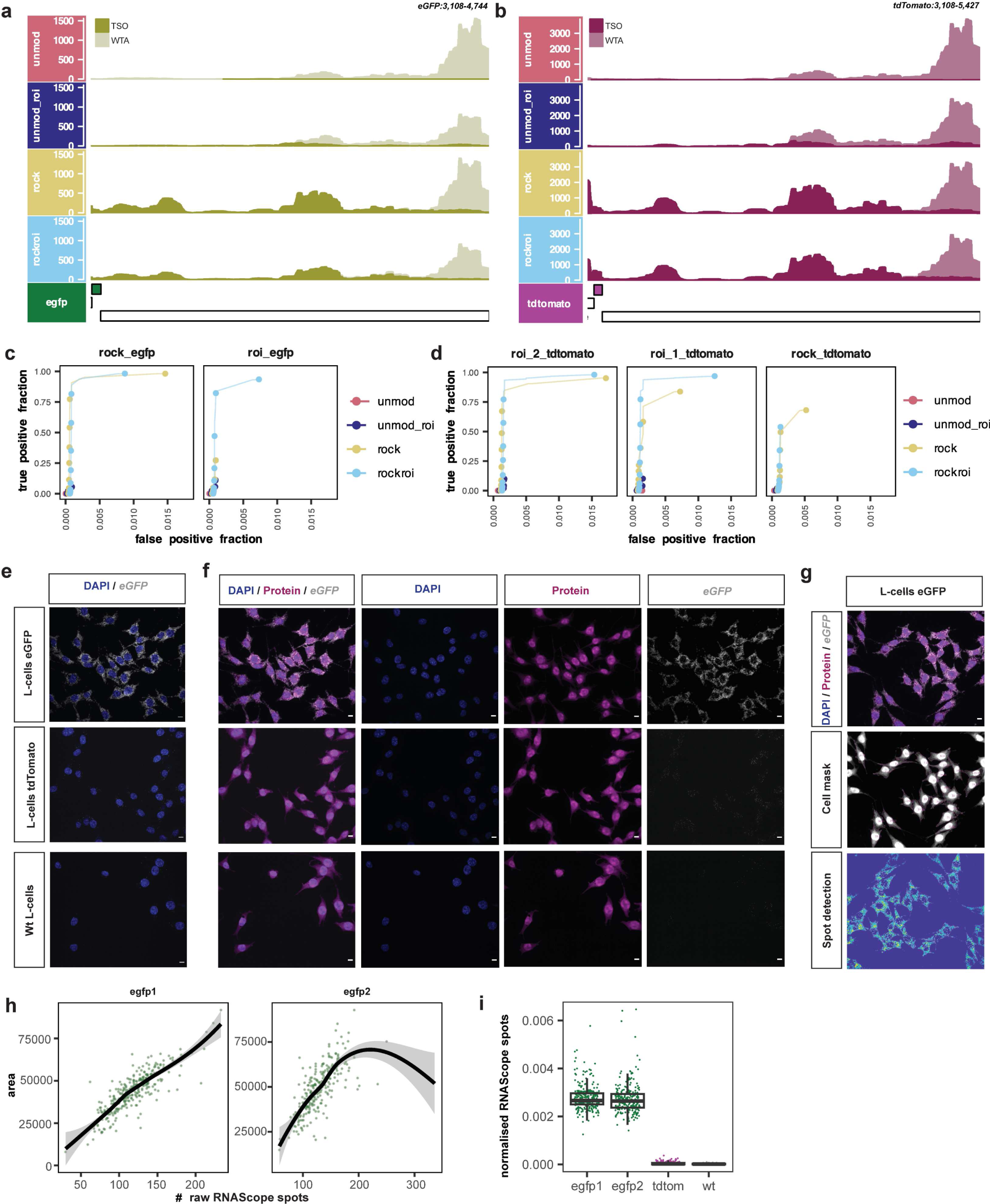
Analysis of target data from scRNAseq experiment using mix of cell lines to test RoCK and ROI performance and effect of cell area on quantification of *eGFP* mRNAs. **a-b,** Zoom in of 3’ UTR from coverage plot in Figure 4b **(a)** and Figure 4c **(b). c-d,** Receiver operating characteristic (ROC) curves indicating the true positive and false positive detection of RoCKseq and ROIseq regions for *eGFP* **(c)** and *tdTomato* **(d).** True positive fraction: detection of *eGFP* in mouse cells or *tdTomato* in human cells, respectively. False positive: detection of *eGFP* in human cells or *tdTomato* in mouse cells, respectively. **e-f,** Detection of *eGFP* transcript in L-cells expressing eGFP, tdTomato or wt L-cells (untransduced) without **(e)** or with **(f)** protein stain. Scale bars: 10 µm. **g,** Example of cell mask and spot detection on L-cells expressing eGFP. Scale bars: 10 µm. **h,** Number of RNAScope spots versus cell area for the two replicates (egfp1 and egfp2). **i,** Number of RNAScope spots normalized by cell area.

**Supp Figure 8:**
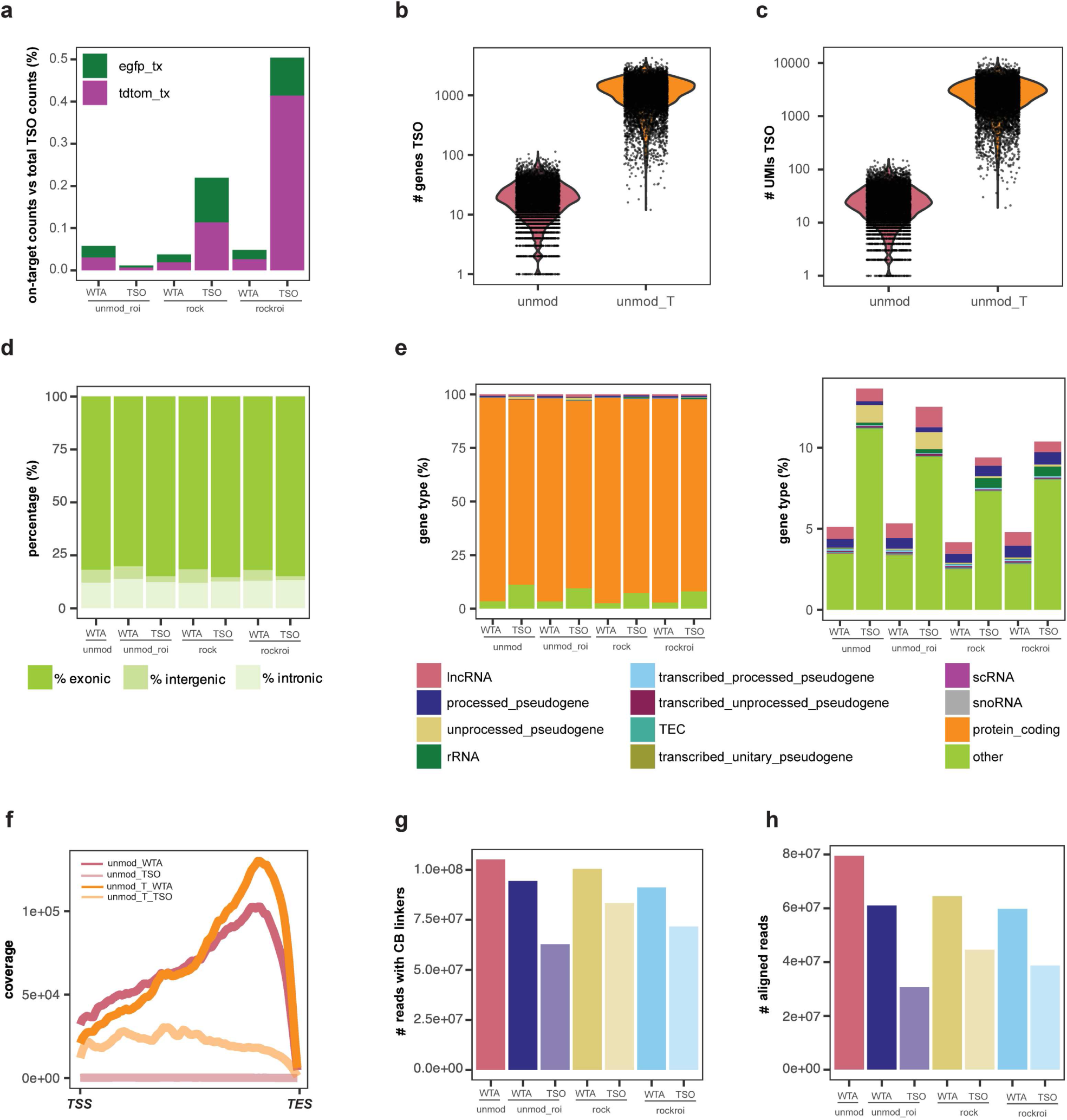
Characterization of TSO data from RoCK and ROI experiments. **a,** On-target counts versus total TSO counts for *eGFP* and *tdTomato* across conditions, including CDS and UTRs. **b-c,** Number of genes **(b)** and UMIs **(c)** detected in the TSO data**. d,** Percent exonic, intergenic and intronic alignments in WTA and TSO data. **e,** Top 11 gene biotypes detected in TSO and WTA data with (right) and without (left) protein coding genes **(e).** Other: all other gene types not in the top 11. **f,** Aggregated gene body sequencing coverage along all transcripts detected in TSO and WTA TSS: transcription start site, TES: transcription end site. **g,** Number of raw reads with canonical WTA and TSO cell barcodes, regardless of whitelists. **h,** Number of aligned reads **(h)**. Numbers indicate the total number of alignments. Data in panels **(a, d-e, g-h)** refer to experiment described in Figure 3 **(a)**, data in panels **(b-c, f)** refer to experiment described in Figure 2 **(b)**.

**Supp Figure 9:**
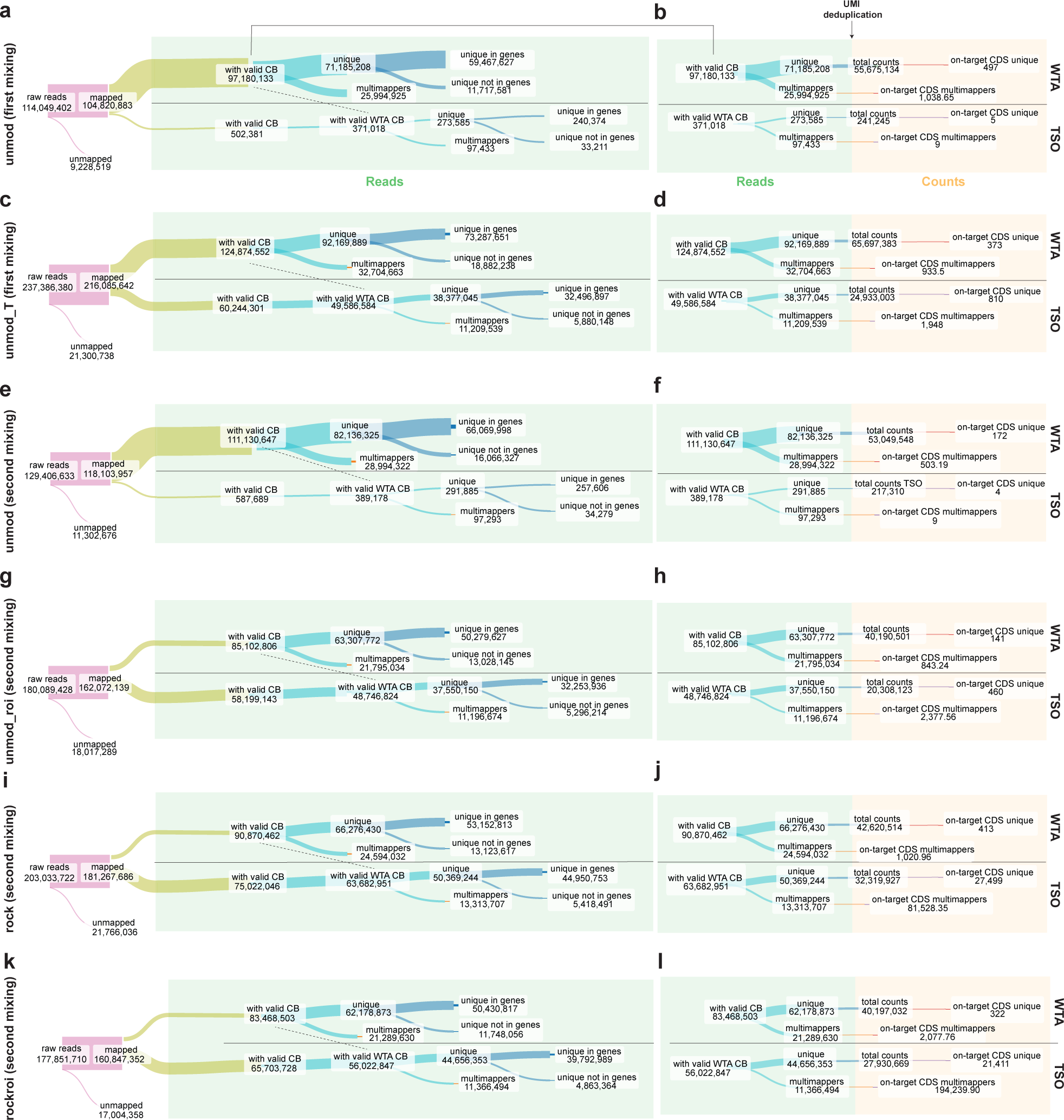
Flow of reads during RoCK and ROI scRNAseq human and mouse mixing experiments. **a-l,** Sankey plots depicting the WTA and TSO sequencing, alignment, UMI deduplication and cell barcode detection performance across conditions (mouse and human mixing experiments). Conditions: unmod (first mixing) and unmod_T (first mixing) for experiment described in Figure 2 **(b) (panels a-d)**; unmod (second mixing), unmod_roi (second mixing), rock (second mixing) and rockroi (second mixing) for experiment described in Figure 3 **(a) (panels e-l)**. Dashed line: filtering of TSO reads with non-empty cells with valid cell barcodes as detected in WTA data. Nodes: <u>raw reads</u>: number of reads from FASTQ files; <u>mapped and unmapped</u>: reads mapped to genome or not; <u>with valid CB (WTA)</u>: WTA reads after EmptyDrop-filtering of cell barcodes (CB) from empty wells; <u>with valid CB (TSO)</u>: TSO reads with a valid cell barcode structure; <u>with valid WTA CB (TSO):</u> TSO reads with cell barcodes matching WTA’s EmptyDrops-filtered cells; <u>unique and multimappers:</u> uniquely and multimapping reads, respectively; <u>unique in genes and unique not in genes</u>: uniquely mapped reads overlapping genes or outside genes, respectively; <u>total counts:</u> total number of (gene) counts after UMI deduplication; <u>on-target CDS unique and on-target CDS multimappers:</u> number of *eGFP* and *tdTomato* counts in their CDS according to the multimapping status of the original read. Counts are 1/n transformed, n being the number of compatible loci (n=1 for unique reads). CDS: coding region.

**Supp Figure 10:**
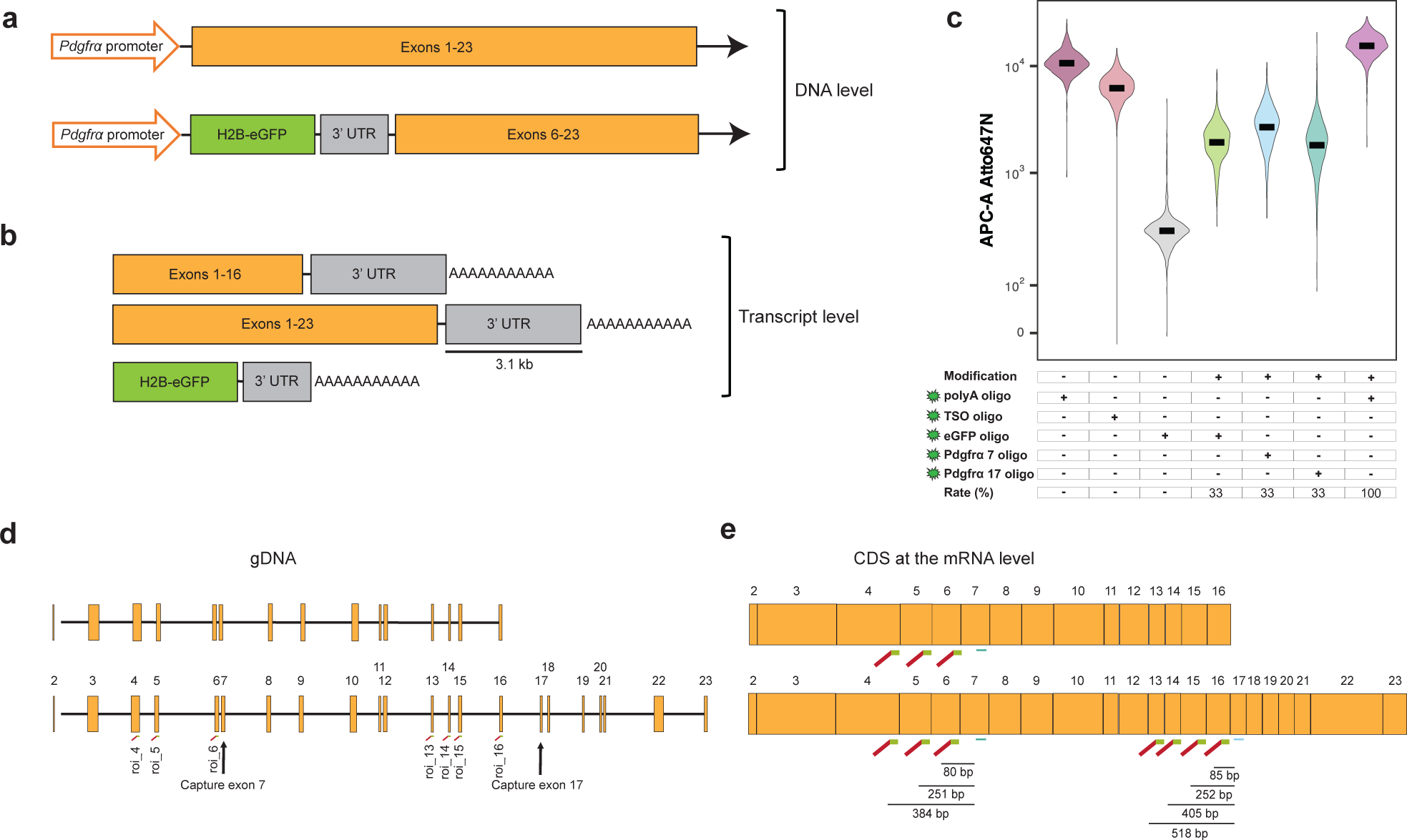
*Pdgfrα* locus, capture, regions of interest and bead modification. **a,** Structure of the *Pdgfrα* locus in the transgenic mouse strain used for the scRNAseq experiment. The mouse strain harbors a *Pdgfrα* allele where the first four exons were substituted with an *H2B-eGFP* construct^52^. **b,** Structure of the *Pdgfrα-*derived transcripts from **(a).** Top two diagrams: long and short *Pdgfrα* isoforms, last diagram: transcript derived from *H2B-eGFP* transgenic allele. **c,** FACS signal from modification of beads for scRNAseq experiment. Y-axis: Atto647N fluorescent signal. The Y-axis has a biexponential transformation. **d-e,** Exons targeted via RoCKseq captures and ROIseq primers for the short (top, ENSMUST00000202681.3 and ENSMUST00000201711.3) and long (bottom, ENSMUST00000000476.14 and ENSMUST00000168162.4) *Pdgfrα* isoforms at the gDNA **(d)** and mRNA **(e)** levels.

**Supp Figure 11:**
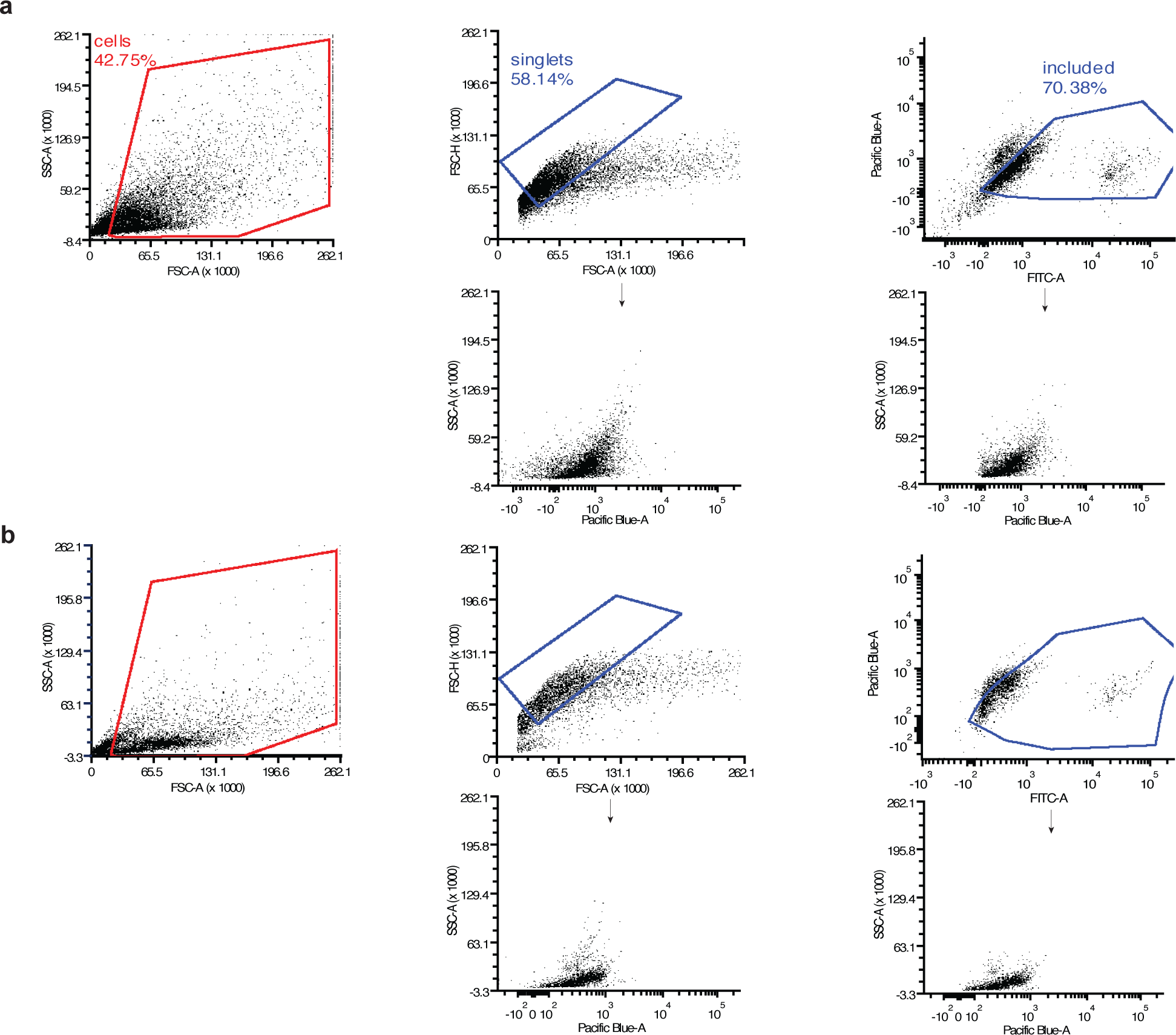
cell gating for murine colonic cells. **a,** FACS gating of the mesenchymal fraction. Gating of cells was done on FSC-A versus SCC-A signal, while singlets were gated in FSC-A versus FSC-H signal. Live cells (included gate) were gated on FITC-A (signal from eGFP positive cells) versus Pacific Blue-A (viability signal). Bottom: additional plots (Pacific Blue-A for live cells versus SSC-A) showing gating for live cells (left: gated for singlets, right: gated for included). **b,** FACS analysis of negative control without viability staining. Gating of cells was done on FSC-A versus SCC-A signal, while singlets were gated in FSC-A versus FSC-H signal. Arrows connect plots that have the same gating but are represented in different channels.

**Supp Figure 12:**
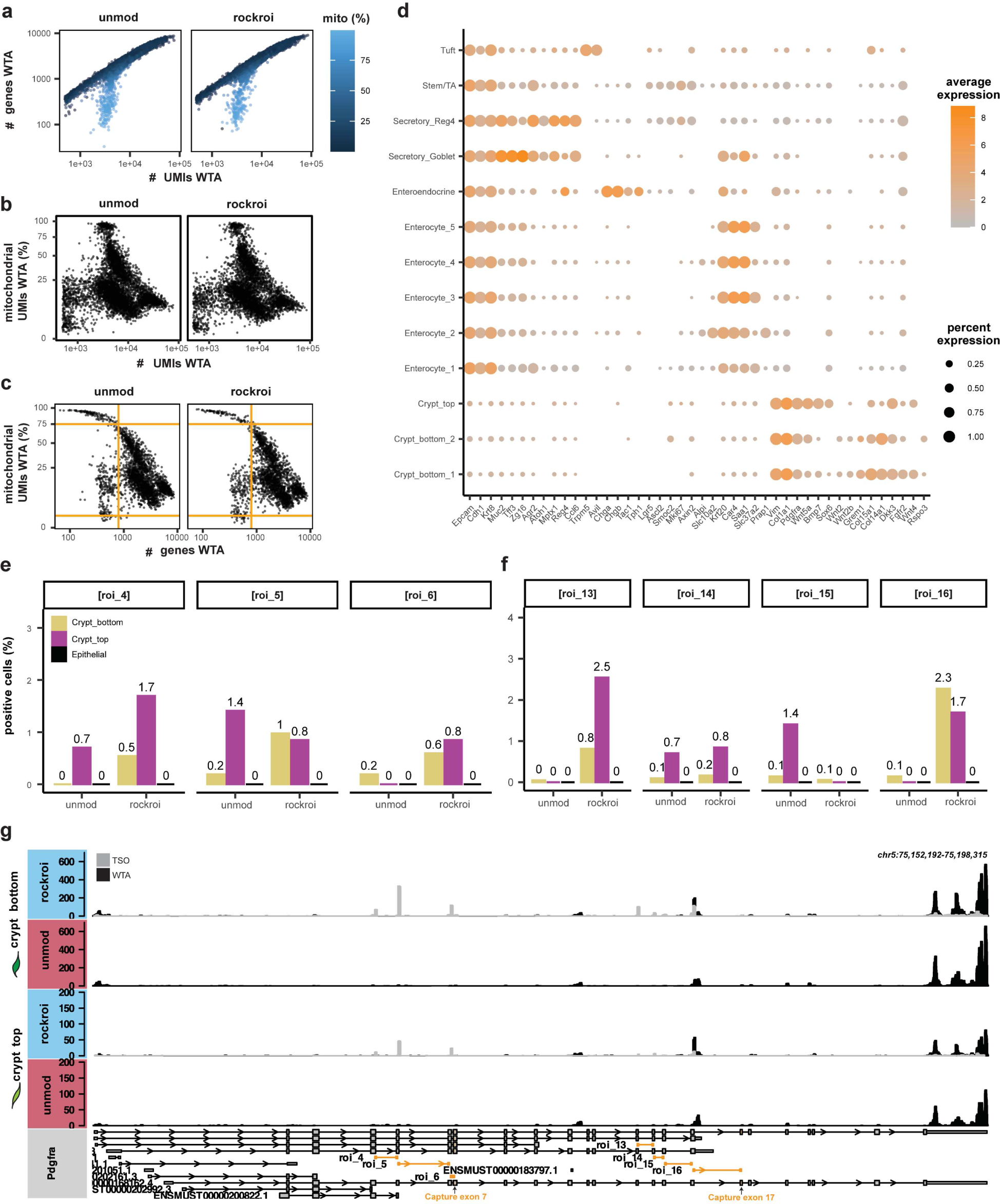
Quality control of *Pdgfrα* scRNAseq experiment, descriptive analysis and splicing quantification. **a,** Number of genes versus number of UMIs colored by mitochondrial content, downsampled WTA data. **b,** Mitochondrial content versus number of UMIs, downsampled WTA data**. c,** Mitochondrial content versus number of genes. Orange lines: QC filtering thresholds. **d,** Expression of manual cell type annotation markers across cell clusters**. e-f,** Percent positive cells in which at least one UMI spanning the splice junction targeted by ROIseq was detected on WTA data. **g,** Coverage along *Pdgfrα* split by crypt top and crypt bottom fibroblasts for TSO (gray) and WTA (black) libraries.

**Supp Figure 13:**
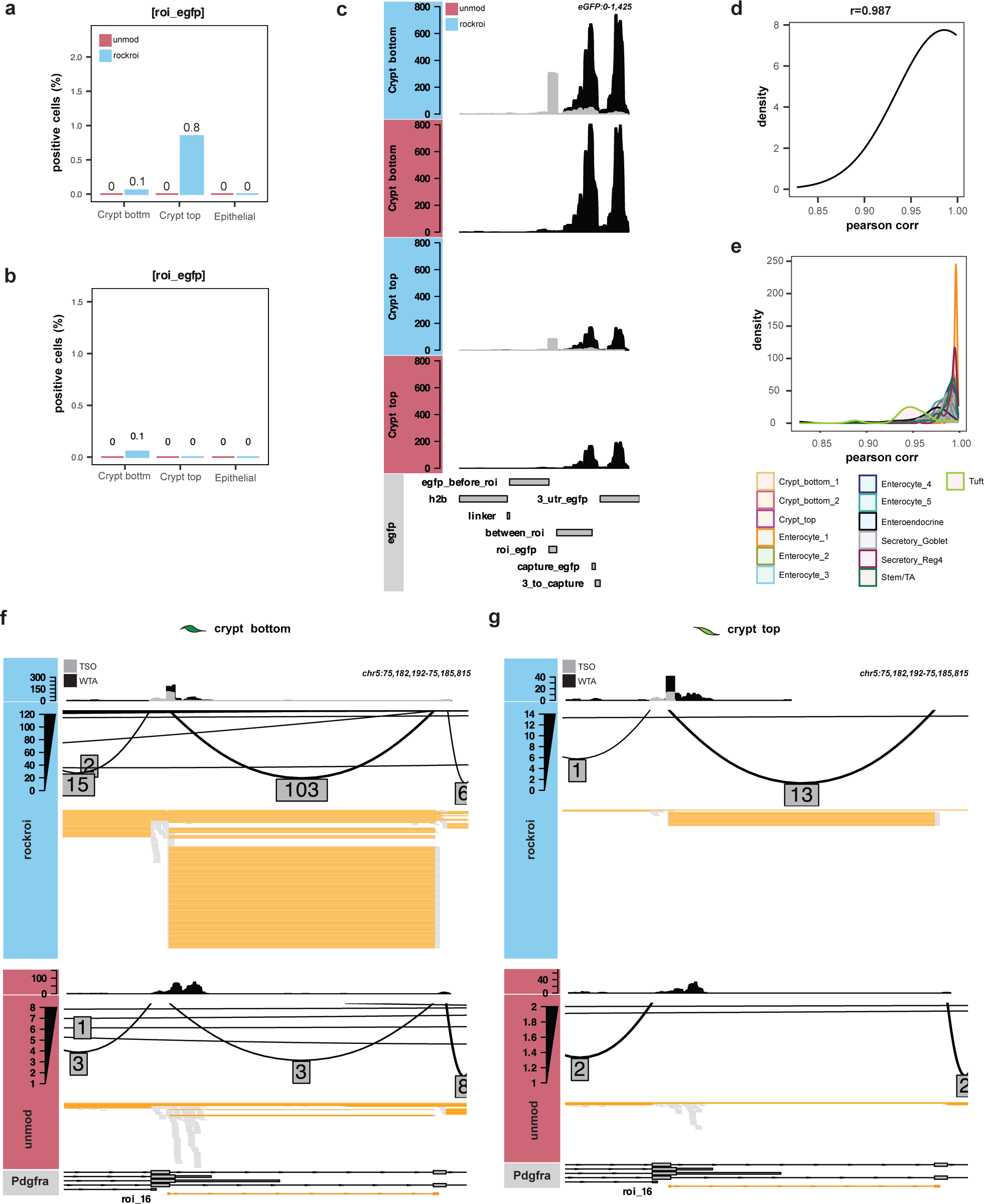
Detection of *Pdgfrα* alternative splicing. **a-b,** Percent positive cells in which at least one UMI for the *eGFP* ROI was detected on TSO **(a)** or WTA data **(b). c,** Coverage along e*GFP***. d-e,** Pearson correlation distributions between same barcodes in the unimodal and multimodal rockroi WTA data for all cells **(d)** or split by cell type **(e**). Correlations were calculated on 100 genes**. f-g,** Coverage, sashimi and alignment tracks for roi_16 region in crypt bottom **(f)** or crypt top **(g)** fibroblasts. Boxed values indicate the number of alignments spanning splice junctions.

## Supplementary tables

**Supp Table 1:** scRNAseq experiments with relevant metrics including sequencing depth and number of cells before and after filtering.

| name_experiment | name_fastqs (R[1,2]) | SRR_identifier | conditions | modalities | bead_modification | number_of_ROIseq_primers | number_of_reads_per_sample | number_of_cells_per_condition_(unfiltered) | number_of_cells_per_condition_(after_QC_filtering_and_doublet_removal) |
| --- | --- | --- | --- | --- | --- | --- | --- | --- | --- |
| First cell line mixing experiment | o307161_1-Unmodified_S4_R*_001.fastq.gz | SRR28817193 | unmod | WTA | No | 0 | 114049402 | 9768 | 8587 |
| First cell line mixing experiment | o307161_2-Unmodified_N1_S1_R*_001.fastq.gz | SRR28817192 | unmod_T | WTA / TSO | No | 0 | 237386380 | 10091 | 7966 |
| Second cell line mixing experiment | 315641_1-Unmod_S4_R*_001.fastq.gz | SRR28830618 | unmod | WTA | No | 0 | 129406633 | 8020 | 6775 |
| Second cell line mixing experiment | 315641_2-Unmod_ROI_S2_R*_001.fastq.gz | SRR28830617 | unmod_roi | WTA / TSO | No | 3 | 180089428 | 7003 | 5724 |
| Second cell line mixing experiment | 315641_3-RoCK_S1_R*_001.fastq.gz | SRR28830616 | rock | WTA / TSO | Yes | 0 | 203033722 | 8226 | 7118 |
| Second cell line mixing experiment | 315641_4-RoCK_ROI_S3_R*_001.fastq.gz | SRR28830615 | rockroi | WTA / TSO | Yes | 3 | 177851710 | 6476 | 5231 |
| Pdgfra epithelial mesenchymal cells | 325411_1-Unmod_WTA_S1_R*_001.fastq.gz | SRR28839349 | unmod | WTA | No | 0 | 379866683 | 5711 | 4634 |
| Pdgfra epithelial mesenchymal cells | 325411_2-RoCK_ROI_WTA_S2_R*_001.fastq.gz | SRR28839348 | rockroi_unimodal | WTA | Yes | 8 | 368399364 | 4980 | 4275 |
| Pdgfra epithelial mesenchymal cells | 325402_1-RoCK_ROI_WTA_S1_R*_001.fastq.gz | SRR28839350 | rockroi_multimodal | WTA / TSO | Yes | 8 | 665200527 | 5079 | 4097 |

**Supp Table 2:** Markers used for annotation of murine colonic cell clusters.

| **Broad_cell_type** | **Cell_type** | **Marker** |
| --- | --- | --- |
| Epithelial | Epithelial | ENSMUSG00000045394.9__Epcam |
| Epithelial | Epithelial | ENSMUSG00000000303.12__Cdh1 |
| Epithelial | Epithelial | ENSMUSG00000049382.10__Krt8 |
| Epithelial | Secretory_cells_Goblet_Reg4 | ENSMUSG00000025515.15__Muc2 |
| Epithelial | Secretory_cells_Goblet_Reg4 | ENSMUSG00000024029.4__Tff3 |
| Epithelial | Secretory_cells_Goblet | ENSMUSG00000049350.6__Zg16 |
| Epithelial | Secretory_cells_Goblet_Reg4 | ENSMUSG00000020581.11__Agr2 |
| Epithelial | Secretory_cells_Reg4 | ENSMUSG00000073043.5__Atoh1 |
| Epithelial | Secretory_cells_Reg4 | ENSMUSG00000026531.4__Mptx1 |
| Epithelial | Secretory_cells_Reg4 | ENSMUSG00000027876.4__Reg4 |
| Epithelial | Secretory_cells_Reg4 | ENSMUSG00000018927.3__Ccl6 |
| Epithelial | Tuft | ENSMUSG00000009246.14__Trpm5 |
| Epithelial | Tuft | ENSMUSG00000025432.11__Avil |
| Epithelial | Enteroendocrine | ENSMUSG00000021194.6__Chga |
| Epithelial | Enteroendocrine | ENSMUSG00000027350.8__Chgb |
| Epithelial | Enteroendocrine | ENSMUSG00000061762.12__Tac1 |
| Epithelial | Enteroendocrine | ENSMUSG00000040046.14__Tph1 |
| Epithelial | Stem_cells | ENSMUSG00000020140.15__Lgr5 |
| Epithelial | Stem_cells | ENSMUSG00000009248.6__Ascl2 |
| Epithelial | Stem_cells | ENSMUSG00000023886.10__Smoc2 |
| Epithelial | Proliferating_stem_cells_and_transiently_amplifying | ENSMUSG00000031004.8__Mki67 |
| Epithelial | Proliferating_stem_cells_and_transiently_amplifying | ENSMUSG00000000142.15__Axin2 |
| Epithelial | Enterocytes | ENSMUSG00000079440.2__Alpi |
| Epithelial | Enterocytes | ENSMUSG00000023073.2__Slc10a2 |
| Epithelial | Enterocytes | ENSMUSG00000035775.2__Krt20 |
| Epithelial | Enterocytes | ENSMUSG00000000805.18__Car4 |
| Epithelial | Enterocytes | ENSMUSG00000074115.5__Saa1 |
| Epithelial | Enterocytes | ENSMUSG00000032122.15__Slc37a2 |
| Epithelial | Enterocytes | ENSMUSG00000025467.8__Prap1 |
| Mesenchymal | Mesenchymal | ENSMUSG00000026728.9__Vim |
| Mesenchymal | Mesenchymal | ENSMUSG00000001506.10__Col1a1 |
| Mesenchymal | Mesenchymal | ENSMUSG00000029231.15__Pdgfra |
| Mesenchymal | Crypt_top_fibroblasts | ENSMUSG00000021994.15__Wnt5a |
| Mesenchymal | Crypt_top_fibroblasts | ENSMUSG00000008999.7__Bmp7 |
| Mesenchymal | Crypt_top_fibroblasts | ENSMUSG00000051910.13__Sox6 |
| Mesenchymal | Crypt_bottom_fibroblast_1_and_2 | ENSMUSG00000010797.6__Wnt2 |
| Mesenchymal | Crypt_bottom_fibroblast_1_and_2 | ENSMUSG00000027840.5__Wnt2b |
| Mesenchymal | Crypt_bottom_fibroblast_1_and_2 | ENSMUSG00000074934.3__Grem1 |
| Mesenchymal | Crypt_bottom_fibroblast_1_and_2 | ENSMUSG00000028339.17__Col15a1 |
| Mesenchymal | Crypt_bottom_fibroblast_1_and_2 | ENSMUSG00000022371.16__Col14a1 |
| Mesenchymal | Crypt_bottom_fibroblast_1_and_2 | ENSMUSG00000030772.6__Dkk3 |
| Mesenchymal | Crypt_bottom_fibroblast_1 | ENSMUSG00000030849.18__Fgfr2 |
| Mesenchymal | Crypt_bottom_fibroblast_1 | ENSMUSG00000036856.4__Wnt4 |
| Mesenchymal | Crypt_bottom_fibroblast_2 | ENSMUSG00000019880.10__Rspo3 |

**Supp Table 3:**
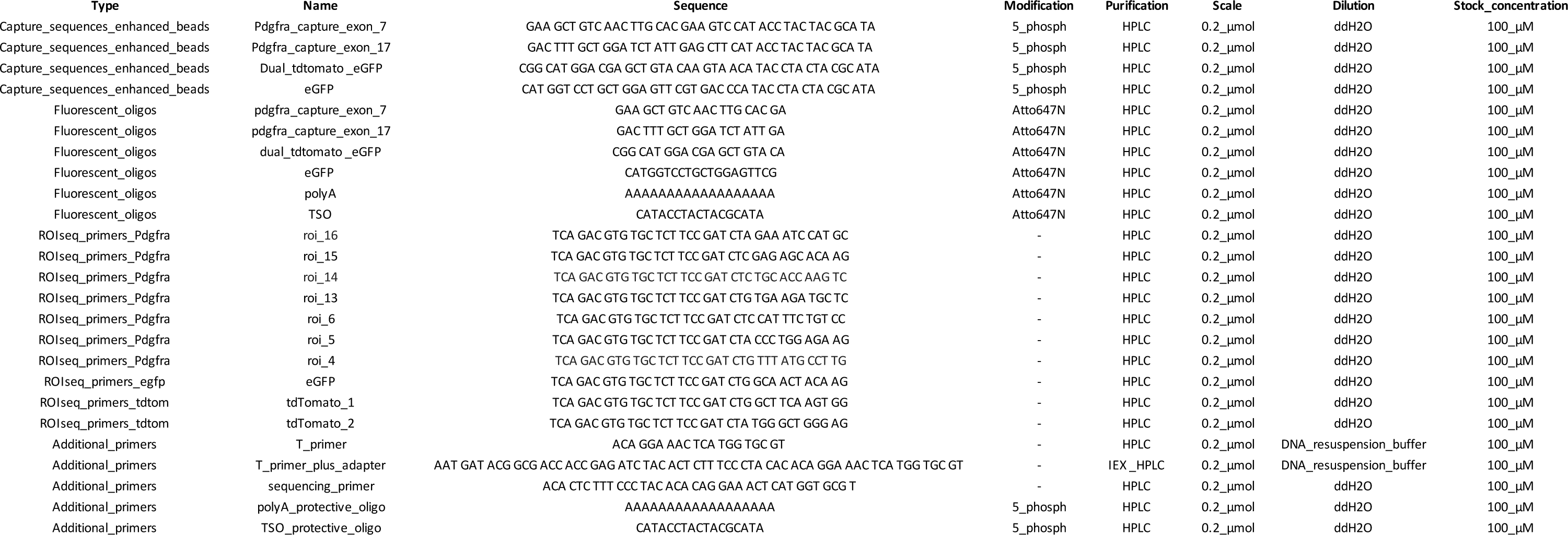
Primer sequences.

